# Antipsychotic-like effects of the selective Rho-kinase 2 inhibitor KD025 in genetic and pharmacological mouse models of schizophrenia

**DOI:** 10.1101/2024.09.16.613372

**Authors:** Rinako Tanaka, Jingzhu Liao, Yue Liu, Wenjun Zhu, Kisa Fukuzawa, Masamichi Kondo, Masahito Sawahata, Daisuke Mori, Akihiro Mouri, Hisayoshi Kubota, Daiki Tachibana, Yohei Kobayashi, Tetsuo Matsuzaki, Taku Nagai, Toshitaka Nabeshima, Kozo Kaibuchi, Norio Ozaki, Hiroyuki Mizoguchi, Kiyofumi Yamada

**Affiliations:** Department of Neuropsychopharmacology and Hospital Pharmacy, Nagoya University Graduate School of Medicine, Nagoya, Aichi, 466-8560, Japan; Department of Applied Pharmacology, Graduate School of Medicine and Pharmaceutical Sciences, University of Toyama, Sugitani, Toyama, 930-0194, Japan; Pathophysiology of mental disorders, Nagoya University Graduate School of Medicine, Nagoya, Aichi, 466-8560, Japan; Brain and Mind Research Center, Nagoya University, Nagoya, Aichi, 466-8560, Japan; Department of Regulatory Science for Evaluation & Development of Pharmaceuticals & Devices, Fujita Health University Graduate School of Medical Science, Toyoake, Aichi, 470-1129, Japan; Division of Behavioral Neuropharmacology, International Center for Brain Science (ICBS), Fujita Health University, Toyoake, Aichi, 470-1192, Japan; Pharmacology Research Unit, Sumitomo Pharma Co., Ltd., Osaka, Osaka, 554-0022, Japan; Laboratory of Health and Medical Science Innovation, Fujita Health University Graduate School of Medical Sciences, Toyoake, Aichi, 470-1192, Japan; Japanese Drug Organization of Appropriate Use and Research, Nagoya, Aichi, 468-0069, Japan; Division of Cell Biology, ICBS, Fujita Health University, Toyoake, Aichi, 470-1129, Japan; Institute for Glyco-core Research (iGCORE), Nagoya University, Furo-cho, Chikusa-ku, Nagoya, Aichi, 464-8601, Japan; ICBS, Fujita Health University, Toyoake, Aichi, 470-1129, Japan

## Abstract

Copy number variations in the *ARHGAP10* gene encoding Rho GTPase–activating protein 10 are significantly associated with schizophrenia. ARHGAP10 negatively regulates RhoA/Rho-kinase (ROCK) signaling. We previously demonstrated that fasudil, a non-selective ROCK inhibitor, exhibited antipsychotic-like effects in several mouse models of schizophrenia. ROCK has two subtypes, ROCK1 and ROCK2. ROCK1 is mainly expressed in the thymus and blood, while ROCK2 is predominantly expressed in the brain. Therefore, it is expected that like fasudil, selective ROCK2 inhibitors will exhibit antipsychotic-like effects, accompanied by a lower incidence of adverse effects due to ROCK1 inhibition. Here, we used genetic and pharmacological models of schizophrenia to investigate whether the selective ROCK2 inhibitor KD025 would show antipsychotic-like effects with a favorable adverse effect profile. Oral administration of KD025 suppressed the abnormal increase in the phosphorylation level of myosin phosphatase–targeting subunit 1, a substrate of ROCK, and ameliorated the decreased spine density of layer 2/3 pyramidal neurons in the medial prefrontal cortex of *Arhgap10* S490P/NHEJ mice. Furthermore, KD025 mitigated the methamphetamine-induced impairment of visual discrimination (VD) in *Arhgap10* S490P/NHEJ and wild-type mice. KD025 also reduced MK-801–induced impairments of VD, novel object recognition, and hyperlocomotion. Regarding side effects that are commonly seen with typical antipsychotics, KD025 did not affect systolic blood pressure and did not induce extrapyramidal symptoms, hyperprolactinemia, or hyperglycemia at the effective dosage in naïve wild-type mice. Taken together, KD025 shows antipsychotic-like effects with a favorable adverse effect profile in genetic and pharmacological mouse models of schizophrenia.

## 2 Introduction

Schizophrenia is a debilitating disorder characterized by positive symptoms, negative symptoms, and cognitive dysfunction^1, 2^. Current pharmacological treatments for schizophrenia have numerous shortcomings in terms of efficacy^3^. First-generation antipsychotics (FGAs) such as haloperidol competitively block the dopamine D2 receptor (D2R) and reduce positive symptoms, but have only a minimal effect on negative symptoms and cognitive dysfunction^4^. In addition, FGAs exhibit severe adverse effects, including extrapyramidal symptoms (EPSs), hyperprolactinemia, and sedation, often resulting in treatment discontinuation^5, 6^. To overcome these issues, second-generation antipsychotics (SGAs), such as risperidone and olanzapine, were developed to block multiple neurotransmitter receptors in addition to D2Rs, but their efficacy against negative symptoms and cognitive dysfunction is still limited^2, 7, 8^. In addition, they can cause severe adverse effects such as hyperglycemia, weight gain, QTc prolongation, and sedation^2, 5–8^. The response rate of these drugs is about 70%, and clozapine, the only drug indicated for treatment-resistant schizophrenia, improves symptoms in only about 30– 60% of treatment-resistant patients^2, 9–12^. Therefore, there remains an unmet need for pharmacological treatments of schizophrenia that have greater clinical efficacy but fewer adverse effects^1, 5, 8^.

Schizophrenia is caused by genetic risk factors and environmental triggers, the latter including perinatal infection and inflammation, as well as the use of cannabis or methamphetamine (METH)^1, 2, 13^. In a recent genome-wide association study, common variant associations at 287 distinct genomic loci were correlated with schizophrenia and were expressed mainly in excitatory and inhibitory neurons of the central nervous system^14^. In addition, a small number of rare copy number variants (CNVs) with moderate to large effect sizes (odds ratios of 2–60) have been identified in schizophrenia^2^. We previously identified a significant association between schizophrenia and CNVs in the *ARHGAP10* gene, which encodes Rho GTPase–activating protein 10 (odds ratio of 12.3)^15^. ARHGAP10 is a Rho GTPase–activating protein that inactivates RhoA/Rho-kinase (ROCK) signaling by converting the GTP-bound form of RhoA to the GDP-bound form^16, 17^. In addition to *ARHGAP10,* several RhoA-associated variants in GTPase-activating proteins and guanine nucleotide exchange factors have been reported to be significantly associated with schizophrenia^18^. To elucidate the pathophysiological role of *ARHGAP10* gene variants, we have developed a genetic mouse model of schizophrenia (*Arhgap10* S490P / NHEJ (non-homologous end-joining) mice) that mimics the *ARHGAP10* variants identified in schizophrenia patients^15^. *Arhgap10* S490P/NHEJ mice carry a double hit involving two variants: a missense variant (p.S490P) and an exonic deletion caused by NHEJ on the other allele^15^. ARHGAP10 S490P showed a lower affinity for active RhoA^15^; therefore, loss of function of ARHGAP10 induced abnormal activation of RhoA/ROCK signaling in the mPFC and striatum of *Arhgap10* S490P/NHEJ mice^19, 20^. Through comprehensive behavioral, histological, and pharmacological studies, we demonstrated that *Arhgap10* S490P/NHEJ mice exhibited METH-induced cognitive impairments and decreased spine density in the medial prefrontal cortex (mPFC)^15, 20^, and these were mitigated by the ROCK inhibitor fasudil^19^. Furthermore, in pharmacological mouse models of schizophrenia, we showed that fasudil ameliorated both METH- and MK-801–induced schizophrenia-relevant behaviors^21, 22^. Accordingly, we propose that ROCK may be a potential therapeutic target in schizophrenia.

ROCK has two subtypes, ROCK1 and ROCK2, both of which are inhibited by fasudil^23, 24^. ROCK1 is mainly expressed in the thymus and blood, while ROCK2 is predominantly expressed in the brain and heart^23, 25^. Although we have demonstrated antipsychotic-like effects of fasudil in both genetic (*Arhgap10* S490P/NHEJ mice) and pharmacological (METH- or MK-801–treated mice) models, the compound induces hypotension, which may impede its clinical application in schizophrenia^26, 27^. Accordingly, we have focused on the selective ROCK2 inhibitor KD025 (belumosudil), which was approved for chronic graft-versus-host disease (cGVHD) by the United States Food and Drug Administration in 2021^28^. Of note, KD025 is brain permeable and inhibits ROCK in the mouse brain^24^. To verify our original hypothesis that ROCK is a potential therapeutic target in schizophrenia, we investigated the effect of the KD025 on the decreased spine density in the mPFC of *Arhgap10* S490P/NHEJ mice, and on METH- and MK-801–induced schizophrenia-relevant behaviors in *Arhgap10* S490P/NHEJ and wild-type (WT) mice. In addition, we investigated whether KD025 induced the typical adverse effects observed both with a non-selective ROCK inhibitor and with currently used antipsychotics.

## 3 Materials and methods

### 3.1 Animals

*Arhgap10* S490P/NHEJ mice were generated on a C57BL/6J genetic background^15^. Male mice aged over 7 weeks were used in the experiment. Animals were handled in accordance with the guidelines established by the Institutional Animal Care and Use Committee of Nagoya University, the Guiding Principles for the Care and Use of Laboratory Animals approved by the Japanese Pharmacologic Society, and the Guide for the Care and Use of Laboratory Animals by the National Institutes of Health of the United States. Further information on the animals used in this study is provided in the Supplementary Materials.

### 3.2 Drug treatment

KD025 (Slx-2119) (Cat#HY-15307/CS-0776; MedChemExpress, Monmouth Junction, NJ, USA) was suspended in 0.1% carboxymethylcellulose–saline. Fasudil monohydrochloride salt (purity > 99%) was kindly supplied by Asahi Kasei Pharma (Tokyo, Japan). Haloperidol (Mitsubishi Tanabe Pharma Corporation, Osaka, Japan) and clozapine (Tokyo Chemical Industry, Tokyo, Japan) were used as control drugs. METH (0.3 or 1 mg/kg) or MK-801 (0.1–0.3 mg/kg) was administrated intraperitoneally 30 min before the tests. The drug doses and timing were based on previous research^19, 21, 22, 24^. Detailed information concerning the schedule of drug treatment is shown in the Supplementary Materials and Supplementary Table S1.

### 3.3 Immunohistochemistry

Immunohistochemistry was conducted as previously described^19^. Detailed information on the antibodies used in this study is shown in Supplementary Table S2. Fluorescence images were captured using a confocal laser microscope (LSM710; Carl Zeiss AG, Oberkochen, Germany) with a 20×/0.8 NA objective lens, and the number of positive cells was counted blindly using the Fiji/ImageJ software package^29^. Further information on the immunohistochemistry experiments is provided in Supplementary Materials.

### 3.4 Golgi staining

Golgi staining was carried out with the FD Rapid Golgi Stain Kit (FD NeuroTechnologies, Ellicott City, MD, USA) using the methods described in a previous study^19^. We obtained the images of layer 2/3 pyramidal neurons in the mPFC by BZ9000 bright-field microscopy (KEYENCE, Osaka, Japan) using a 100×/1.40 NA oil-immersion objective lens. We quantified the spine density of dendrites 50–200 μm from the soma of pyramidal neurons (six dendrites per mouse) using Neurolucida software and NeuroExplorer (MicroBrightField Bioscience, Williston, VT, USA). Spine density was expressed as the number of spines per 10 μm of dendrite length. Further information on Golgi staining is provided in Supplementary Materials.

### 3.5 Behavioral experiments

To motivate mice to perform a touchscreen-based visual discrimination (VD) task, food and water were restricted to 1 h per day from at least 1 week before pre-training until the end of the task, with the goal of achieving approximately 85–95% of the original bodyweight^19^. A pre-training phase conditioned mice to associate screen contact with the delivery of a liquid reward. Subsequently two stimuli (marble and fan) were presented simultaneously in two different windows, only one of which was associated with the correct response. Touching the correct window (correct response) led to presentation of a liquid reward (20 μL), while touching the incorrect one (incorrect response) resulted in a 5-s time-out period. The percentage of correct responses and the latency from the correct response to reward retrieval (reward latency) were analyzed. The novel object recognition test (NORT)^22^, locomotor test^22^, and open-field test^30^ were performed as previously described. In the NORT, which is used to measure cognitive function, the preference index in the retention session was calculated as the ratio of the amount of time spent exploring the novel object to the total time spent exploring both objects. The pole test^31, 32^ and bar test^33, 34^ were performed as previously described, with minor modifications. In the pole test, the times required for the animal to completely rotate downward from the head-upward position (T_turn_) and descend to the floor (T_total_) were measured, with a maximum limit of 300 s. One mouse with a T_total_ over 300 s before treatment and one that preferred to climb the bar were excluded. Further information on the behavioral experiments is provided in the Supplementary Materials.

### 3.6 Measurement of systolic blood pressure and heart rate

Systolic blood pressure and heart rate were measured using a tail-cuff system (MK 2000; Muromachi Kikai, Tokyo, Japan)^35^ as described in the Supplementary Materials. Considering the variations in measurement times due to differences in the T_max_ of test compounds, we confirmed that blood pressure did not change at all between 20 min and 120 min after vehicle treatment (data not shown).

### 3.7 Measurement of serum prolactin and blood glucose concentrations

Serum prolactin concentrations were assessed with the Mouse Prolactin ELISA Kit (ab100736; Abcam, Cambridge, UK) using an overnight incubation protocol^36, 37^. Blood glucose concentrations of mice fasted for about 18 h were measured in duplicate with the FreeStyle Freedom Lite Blood Glucose Monitoring System (11-779; NIPRO CORPORATION, Osaka, Japan)^38^.

### 3.8 Statistical analysis

The sample size in each experiment was determined with reference to our previous studies according to experiment type^19, 21, 22, 31–38^. Detailed information concerning statistical analysis is shown in the Supplementary Materials and Supplementary Table S3.

## 4 Results

### 4.1 KD025 ameliorated the decrease spine density in mPFC and the impairment in VD task performance induced by a low dose of METH (0.3 mg/kg) in *Arhgap10* S490P/NHEJ mice

To verify the effects of the selective ROCK2 inhibitor KD025, we first confirmed ROCK2 expression in the brain of C57BL/6 mice by immunohistochemistry. ROCK2 signals were detectable in the mPFC and striatum (Supplementary Figure S1A), and predominantly with neuronal nuclei (NeuN)–positive neurons in the mPFC and striatum, but only minimally with Iba-1– or glial fibrillary acidic protein (GFAP)–positive cells (Supplementary Figure S1), suggesting that ROCK2 is mainly expressed in neurons of the mPFC and striatum. Subsequently, we performed immunochemistry analysis to determine whether the systemic administration of KD025 would suppress the phosphorylation level of myosin phosphatase–targeting subunit 1 (MYPT1), a substrate of ROCK, in the brain of *Arhgap10* S490P/NHEJ mice, as seen with fasudil^19^. The ratio of the number of phosphorylated MYPT1 (pMYPT1)–positive cells to that of NeuN-positive cells in the mPFC of *Arhgap10* S490P/NHEJ mice was larger than that in WT mice. KD025 significantly reduced the increased ratio in *Arhgap10* S490P/NHEJ mice, but not that in WT mice (Figure 1A, B). These results suggest that oral administration (p.o.) of KD025 can suppress the abnormal activation of ROCK in mPFC neurons of *Arhgap10* S490P/NHEJ mice.

**Figure 1.**
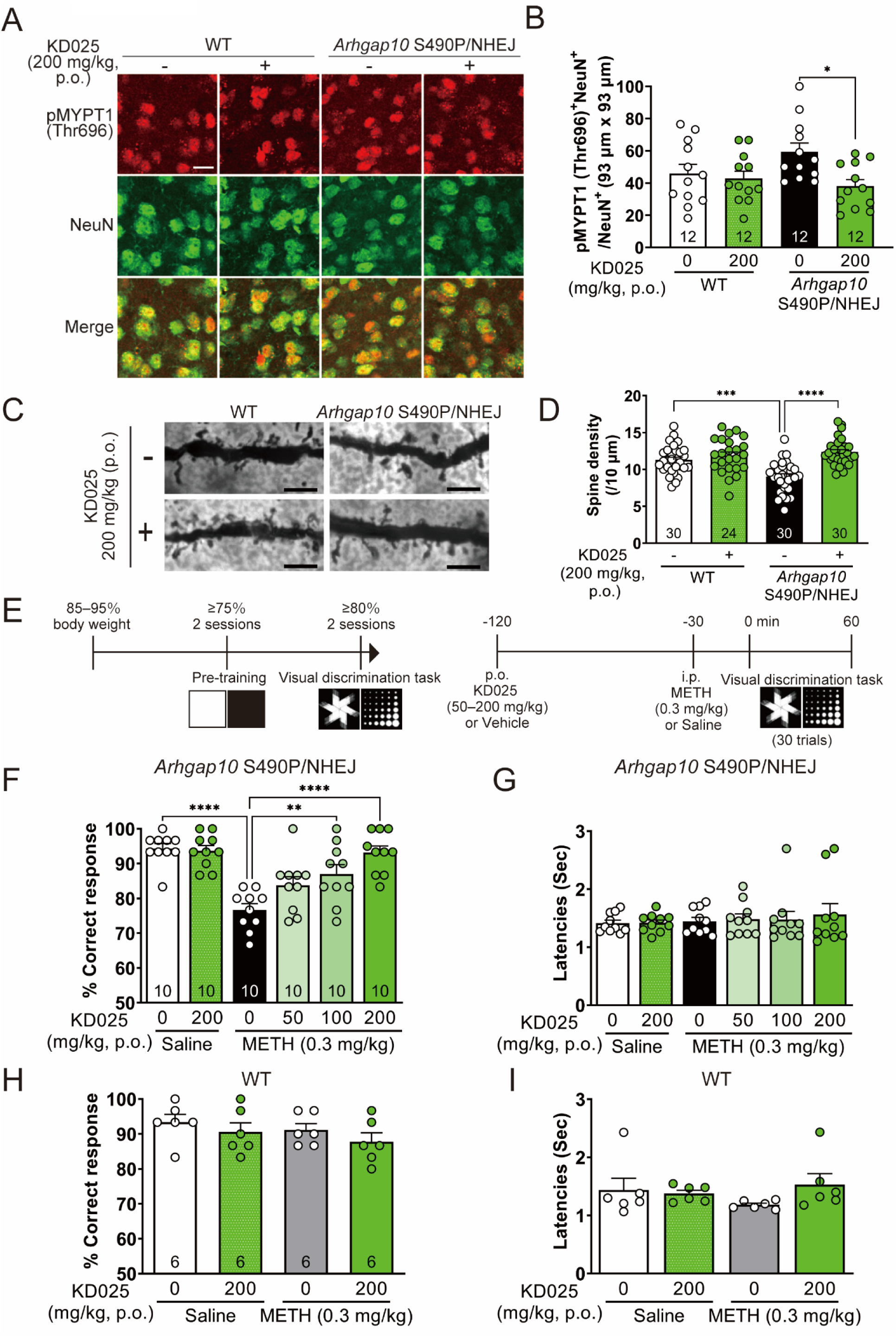
KD025 ameliorated the decreased spine density in the mPFC and the impairment in the VD task induced by a low dose of METH (0.3 mg/kg) in *Arhgap10* S490P/NHEJ mice. A: Representative images of pMYPT1 (Thr696) (red) and NeuN (green) immunoreactivity in the mPFC of *Arhgap10* S490P/NHEJ mice (scale bar indicates 10 μm). B: Ratio of pMYPT1 (Thr696) and NeuN double-positive neurons to NeuN-positive neurons in the mPFC of *Arhgap10* S490P/NHEJ mice or WT mice 120 min after treatment with or without KD025 (200 mg/kg, p.o.). Data represent the mean + SEM (n = 12 slices per group; (four mice per group, three slices per mouse)) and were analyzed by Tukey’s multiple comparison test (*P < 0.05). C: Representative images of dendric spines (scale bar indicates 5 μm). D: Spine density of layer 2/3 pyramidal neurons in the mPFC of *Arhgap10* S490P/NHEJ mice or WT mice with or without KD025 (200 mg/kg, p.o.) treatment for 7 days. Data represent the mean + SEM (n = 24–30 dendrites (four to five mice per group, six dendrites per mouse) and were analyzed by Tukey’s multiple comparison test (***P < 0.001 and ****P < 0.0001). E: The protocols of the VD task (left) and drug treatment (right). F–I: Percentage of correct responses (F) and reward latency (G) in the VD task in *Arhgap10* S490P/NHEJ mice. Percentage of correct responses (H) and reward latency (I) in the VD task in WT mice. The animals were treated with low-dose METH (0.3 mg/kg, i.p.) or KD025 (50–200 mg/kg, p.o.) 30 min or 120 min before the task, respectively. Data represent the mean + SEM (n = 10 for *Arhgap10* S490P/NHEJ mice and n = 6 for WT mice) and were analyzed by Tukey’s multiple comparison test (**P < 0.01 and ****P < 0.0001).

We previously demonstrated that the spine density of layer 2/3 pyramidal neurons in the mPFC was significantly decreased in *Arhgap10* S490P/NHEJ mice compared with WT mice, and this reduction was rescued by fasudil treatment for 7 days^19^. Thus, in our next experiment we administered KD025 (200 mg/kg) once a day for 7 days and showed that this treatment mitigated the decreased spine density of layer 2/3 pyramidal neurons in the mPFC of *Arhgap10* S490P/NHEJ mice, while it did not affect the spine density in WT mice (Figure 1C, D). This result suggests that ROCK2 inhibition is sufficient to ameliorate the decreased spine density in the mPFC of *Arhgap10* S490P/NHEJ mice.

We also previously reported that *Arhgap10* S490P/NHEJ mice showed impairment in the touchscreen-based VD task following a low dose of METH treatment (0.3 mg/kg), whereas it had no effect on cognitive performance in WT mice^20^. This suggests a vulnerability to METH in *Arhgap10* S490P/NHEJ mice, and is consistent with the fact that the symptoms of schizophrenia patients are exacerbated by psychostimulant drugs^39, 40^. Fasudil treatment was shown to mitigate the VD impairment caused by low-dose METH administration in *Arhgap10* S490P/NHEJ mice^19^; therefore, we investigated whether this outcome would also be observed with KD025 treatment. *Arhgap10* S490P/NHEJ mice that exhibited stable discrimination performance (≥ 80% correct responses) for 2 consecutive days were treated with KD025 and low-dose METH on the test day (Figure 1E). The low dose of METH (0.3 mg/kg) significantly decreased the percentage of correct responses, while this impairment was reduced by treatment with KD025 (100–200 mg/kg) in a dose-dependent manner (Figure 1F). However, the administration of KD025 and low-dose METH did not change the reward latency (Figure 1G), suggesting that these drugs do not affect locomotor function or motivational state. By contrast, WT mice treated with the low dose of METH (0.3 mg/kg) and KD025 did not exhibit any changes in the percentage of correct responses, and also showed no change in the reward latency (Figure 1H, I). Of note, KD025 (200 mg/kg) had no effect on the percentage of correct responses in *Arhgap10* S490P/NHEJ or WT mice treated with saline (Figure 1F, H). These results suggest that KD025 reduces the cognitive vulnerability to METH in *Arhgap10* S490P/NHEJ mice.

### 4.2 KD025 ameliorated METH (1 mg/kg)-induced cognitive dysfunction in WT mice

The symptoms of METH psychosis have been shown to closely resemble those of schizophrenia, including the presence of positive symptoms and cognitive impairment^41^. In the present study, we used a pharmacological mouse model of schizophrenia involving METH treatment at a dose of 1 mg/kg, which impairs VD task performance in WT mice that do not have *ARHGAP10* variants. We first investigated whether KD025, like fasudil, would suppress the increased MYPT1 phosphorylation levels in the striatum, because the METH-induced activation of pMYPT1 is most pronounced in the striatum^21^. The ratio of the number of pMYPT1-positive cells to that of NeuN-positive cells in the striatum of WT mice was significantly increased by METH (1 mg/kg) but significantly decreased by KD025 (200 mg/kg, p.o.) (Figure 2A, B). This result suggests that METH (1 mg/kg) activates ROCK in the striatum and KD025 suppresses it. Next, we evaluated the effect of KD025 on METH (1 mg/kg)-induced VD task impairment in WT mice. METH (1 mg/kg) significantly decreased the percentage of correct responses, while KD025 ameliorated this impairment in a dose-dependent manner (Figure 2C). However, neither METH alone nor in combination with KD025 significantly decreased the reward latency, suggesting that these drugs have little effect on motor function or motivation state at the doses used (Figure 2D). We also evaluated the effect of KD025 on METH (1 mg/kg)-induced hyperlocomotion, a positive symptom-like behavior, in WT mice. KD025 (200 or 300 mg/kg, p.o.) had little effect on hyperlocomotion in WT mice, while haloperidol (0.3 mg/kg) significantly suppressed it (Supplementary Figure S2). This result is consistent with that of a previous study showing that fasudil failed to suppress METH (1 mg/kg)-induced hyperlocomotion in WT mice^21^. These observations suggest that in mice, KD025 ameliorates METH-induced cognitive dysfunction but not hyperlocomotion.

**Figure 2.**
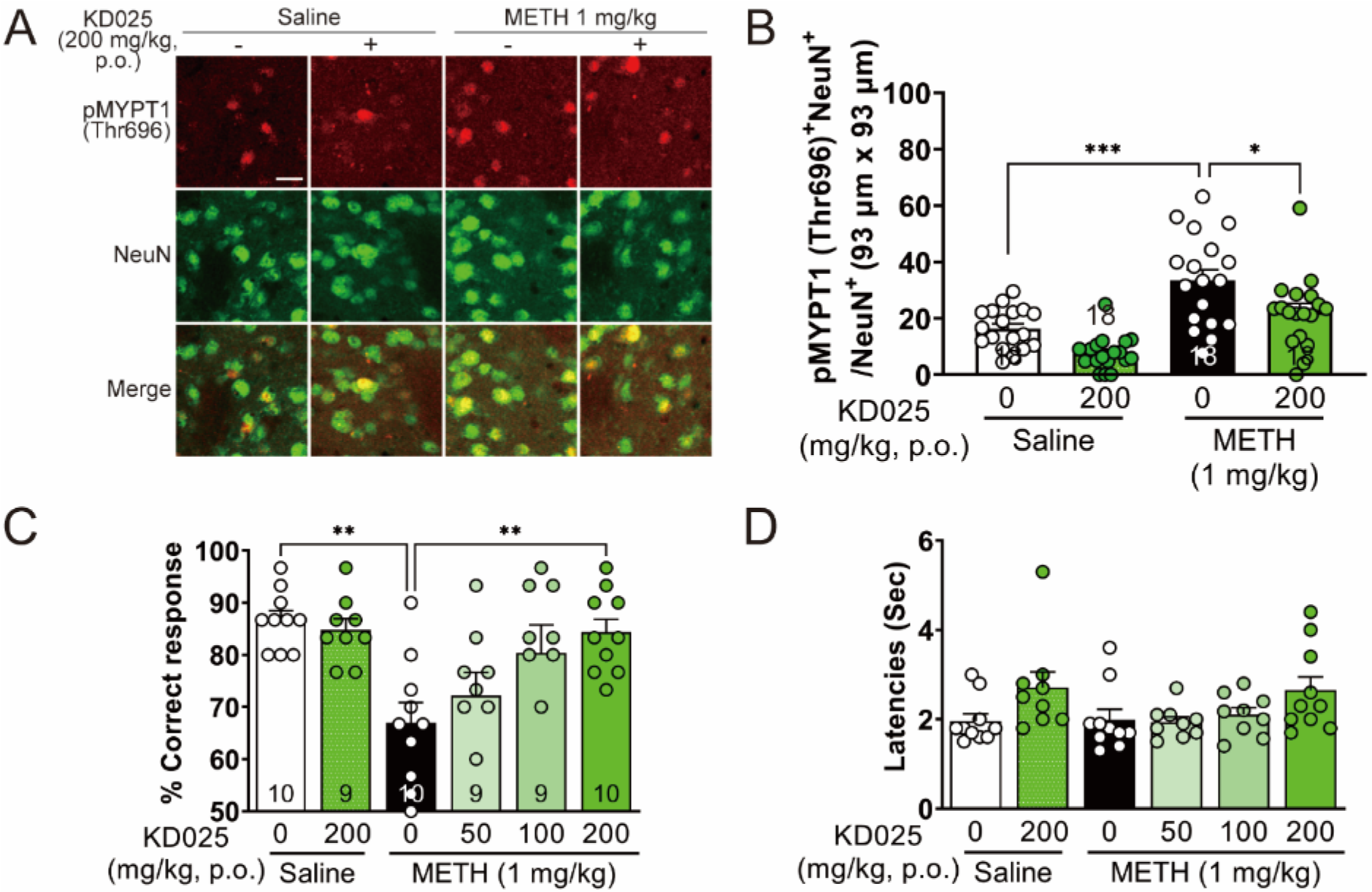
KD025 ameliorated METH (1 mg/kg)-induced cognitive dysfunction in WT mice. A: Representative images of pMYPT1 (Thr696) (red) and NeuN (green) immunoreactivity in the striatum of WT mice (scale bar indicates 10 μm). B: Ratio of pMYPT1 (Thr696) and NeuN double-positive neurons to NeuN-positive neurons in the striatum of WT mice 30 min or 120 min after METH (1 mg/kg, i.p.) or KD025 (200 mg/kg, p.o.) treatment, respectively. Data represent the mean + SEM (n = 18 slices (six mice per group, three slices per mouse)) and were analyzed by Tukey’s multiple comparison test (*P < 0.05, ***P < 0.001). C, D: Percentage of correct responses (C) and reward latency (D) in the VD task in WT mice. The animals were treated with METH (1 mg/kg, i.p.) or KD025 (50– 200 mg/kg, p.o.) 30 min or 120 min before the task, respectively. Data represent the mean + SEM (n = 9–10 WT mice per group) and were analyzed by Tukey’s multiple comparison test (**P < 0.01).

### 4.3 KD025 ameliorated MK-801–induced schizophrenia-relevant behaviors in WT mice

MK-801–induced schizophrenia-relevant behaviors have been utilized in mice as a pharmacological model of schizophrenia on the basis of the glutamate hypothesis^42^. Thus, we evaluated whether KD025 would reduce MK-801–induced impairment in genetic and METH-treatment mouse models of schizophrenia. MK-801 (0.1 mg/kg) significantly decreased the percentage of correct responses in WT mice. KD025 ameliorated MK-801–induced impairment in a dose-dependent manner, without any effects on reward latency (Figure 3A, B). Next we performed the NORT, and found that KD025, like clozapine, mitigated the MK-801–induced decrease in exploratory preference for a novel object in the retention session but not in the training session (Figure 3C, D). Neither MK-801 nor KD025 treatment affected the total exploratory time in the training or retention session (Figure 3E, F). These results suggest that KD025 diminishes the MK-801–induced impairment of object recognition memory in mice. Schizophrenia patients show deficits in sensorimotor gating and impairment of pre-pulse inhibition (PPI)^43^. We previously showed that fasudil partially reduced MK-801–induced impairment of the PPI of the startle response in mice^22^. However, KD025 failed to ameliorate impairment of the PPI in mice (Supplementary Figure S3). These results suggest that KD025 decreases cognitive dysfunction in VD and object recognition memory, but has little effect on sensorimotor gating deficits in the MK-801–treated mouse model of schizophrenia. Lastly, we demonstrated that KD025 (200 mg/kg) significantly suppressed MK-801–induced hyperlocomotion in mice (Figure 3G, H). These results suggest that the selective ROCK2 inhibitor KD025 may have antipsychotic-like effects in schizophrenia.

**Figure 3.**
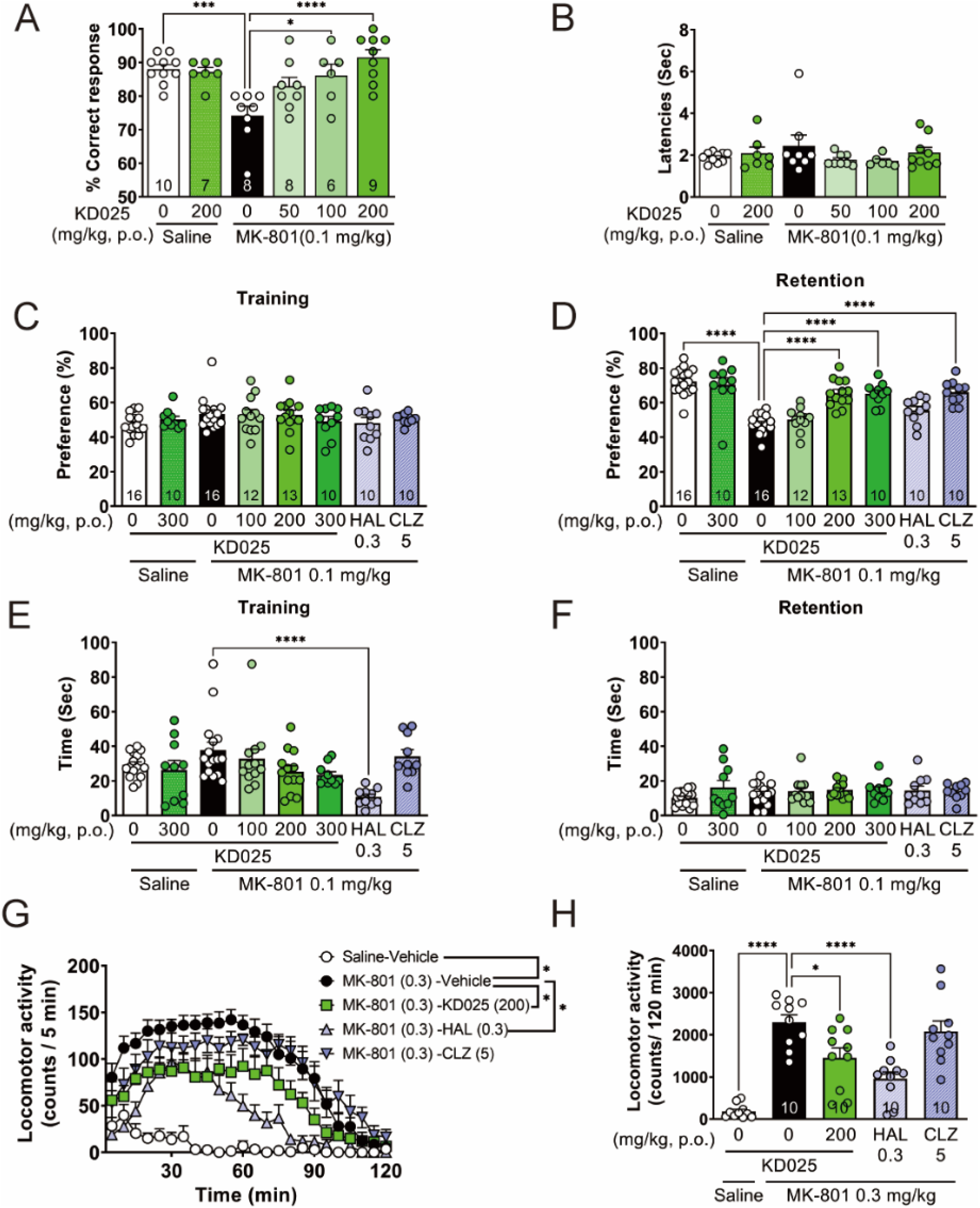
KD025 ameliorated MK-801–induced schizophrenia-relevant behaviors in WT mice. A, B: Percentage of correct responses (A) and reward latency (B) in the VD task in WT mice. The animals were treated with MK-801 (0.1 mg/kg, i.p.) or KD025 (50–200 mg/kg, p.o.) 30 min or 120 min before the task, respectively. Data represent the mean + SEM (n = 6–10 WT mice per group) and were analyzed by Tukey’s multiple comparison test (*P < 0.05, ***P < 0.001, and ****P < 0.0001). C–F: Exploratory preferences in training (C) and retention sessions (D) in the NORT. Total exploration time in the training (E) and retention (F) sessions. MK-801 (0.1 mg/kg, i.p.), haloperidol (0.3 mg/kg, p.o.), clozapine (5 mg/kg, p.o.), or KD025 (100–300 mg/kg, p.o.) was administered 30, 60, 60 or 120 min, respectively, before the training session. The retention session was performed 24 h after the training session. Data are represented as the mean + SEM (n = 10–16 WT mice per group) and were analyzed by Tukey’s multiple comparison test (****P < 0.0001). G–H: Time course changes in locomotor activity (G) and total locomotor activity (H) after MK-801 treatment. Haloperidol (0.3 mg/kg, p.o.), clozapine (5 mg/kg, p.o.), or KD025 (200 mg/kg, p.o.) was administered 60, 60, or 120 min before the test, respectively, and the locomotor activity was measured for 120 min immediately after MK-801 treatment (0.3 mg/kg, i.p.). Data are represented as the mean + SEM (n = 10 WT mice per group) and were analyzed by Tukey’s multiple comparison test (*P < 0.05 and ****P < 0.0001). HAL = haloperidol. CLZ = clozapine.

### 4.4 KD025 did not affect general behavior, locomotion, or blood pressure in naïve WT mice

The aforementioned experiments demonstrated the potential antipsychotic-like effects of KD025 in both genetic and pharmacological mouse models of schizophrenia. Considering the clinical application of selective ROCK2 inhibitors, we investigated whether KD025 would affect general behavior in naïve WT mice. In the open-field test, KD025 (100–300 mg/kg, p.o.) had little effect on any evaluated behavior, including the duration spent in each zone, the total distance moved, and the number of instances of crossing, rearing, jumping, grooming, defecation, and urination (Figure 4A– H). Although fasudil suppressed spontaneous locomotor activity^22^, KD025 (100–300 mg/kg, p.o.) had little effect on this activity during the 180-min post-treatment observation period (Figure 4I). Fasudil induces hypotension in humans and rodents because ROCK mediates arterial smooth muscle contraction^26, 27^. Although fasudil (20 mg/kg, intraperitoneal administration (i.p.) or p.o.) significantly decreased systolic blood pressure compared with vehicle (Figure 4J), KD025 (100–300 mg/kg, p.o.) caused minimal changes in blood pressure and heart rate compared with vehicle (Figure 4J, Supplementary Figure S4). These results suggest that the selective ROCK2 inhibitor KD025 has mild or no effects on general behavior and blood pressure.

**Figure 4.**
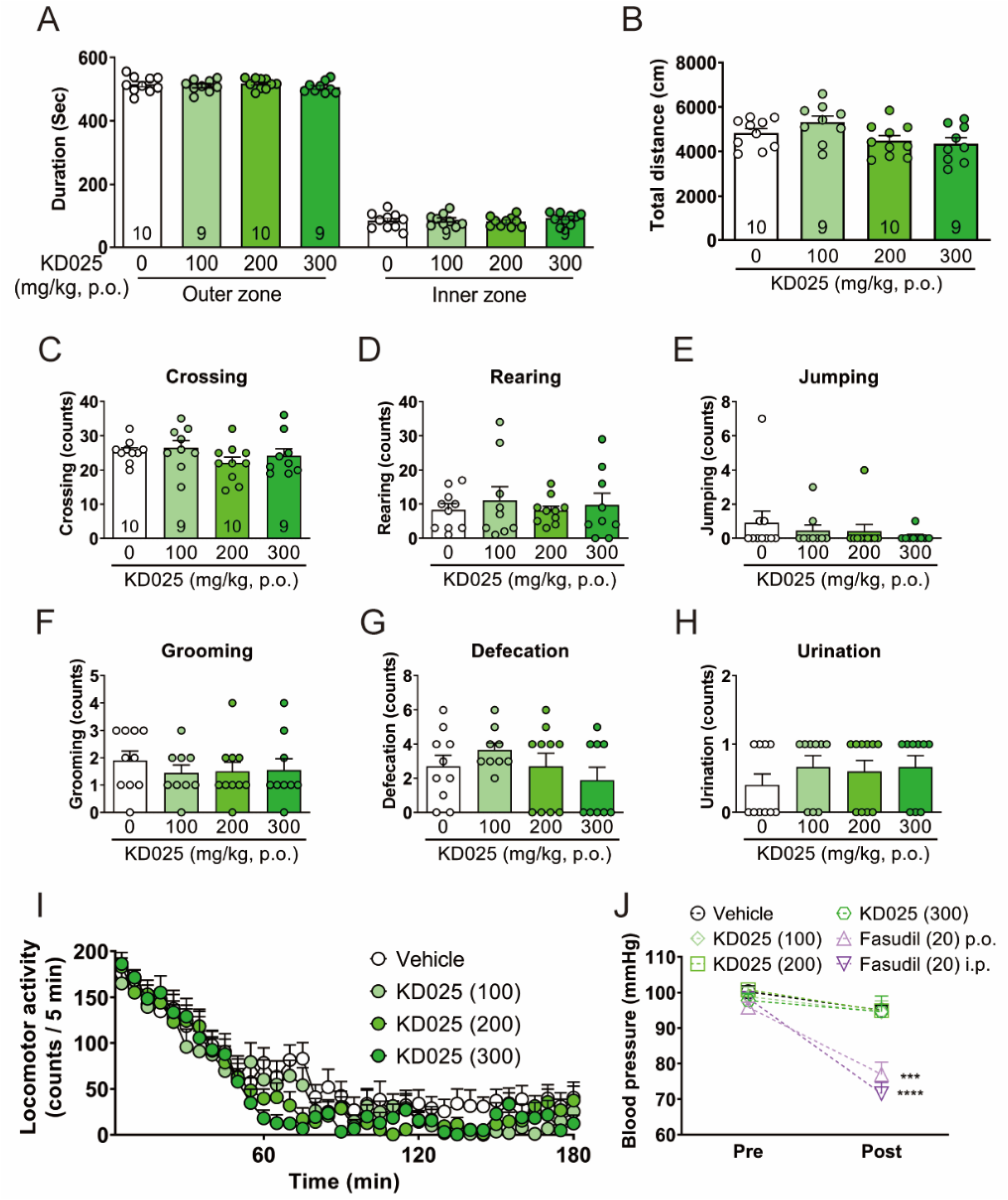
KD025 did not affect general behavior, spontaneous locomotor activity, or blood pressure in naïve WT mice. A–H: The duration in the inner and outer zone (A), the total distance moved (B), and the numbers of instances of crossing between the inner and outer zone (C), rearing (D), jumping (E), grooming (F), defecation (G), and urination (H) for 10 min in the open-field test. KD025 (100–300 mg/kg, p.o.) was administered 120 min before the test. Data represent the mean + SEM (n = 9–10 WT mice per group) and were analyzed by Tukey’s multiple comparison test. I: Time course of locomotor activity after KD025 (100–300 mg/kg, p.o.) treatment. Data are represented as the mean + SEM (n = 9–10 WT mice per group) and were analyzed by Tukey’s multiple comparison test. J: Systolic blood pressure before fasudil (20 mg/kg, i.p. or p.o.) or KD025 (100–300 mg/kg, p.o.) treatment, and 20 min or 120 min afterward, respectively. Data represent the mean + SEM (n = 4–5 WT mice per group) and were analyzed by Šídák’s multiple comparisons test (***P < 0.001, ****P < 0.0001 compared with pre-treatment).

### 4.5 KD025 did not induce EPS or hyperprolactinemia, but increased blood glucose concentrations at high doses in naïve WT mice

Current drugs used to treat schizophrenia, such as D2R antagonists, result in EPS and hyperprolactinemia in clinical use, resulting in significant difficulty with treatment adherence^5, 44^. We therefore examined whether KD025 would induce EPS and hyperprolactinemia. First, we evaluated bradykinesia, a type of EPS characterized by slow voluntary movement^45^. In the pole test, T_total_ was significantly longer 4 h after treatment with haloperidol (2 mg/kg) than vehicle, while KD025 (200 and 1,000 mg/kg) did not change T_total_ or T_turn_ during a 24-h period after treatment (Figure 5A and Supplementary Figure S5). To assess EPS in mice, we also evaluated cataleptogenic behavior. In the bar test, haloperidol (2 mg/kg) increased the time on the bar compared with vehicle during a 6-h period after treatment (Figure 5B). By contrast, KD025 (200, 1,000 mg/kg, p.o.) resulted in comparable times on the bar compared with vehicle during a 24-h period after treatment (Figure 5B). Next, we evaluated serum prolactin concentrations after drug treatment. Haloperidol (0.3 and 2 mg/kg) significantly increased the serum prolactin concentration 1 h after treatment, while KD025 and fasudil had little effect (Figure 5C). Finally, we investigated whether KD025 would increase blood glucose concentrations, because SGAs such as olanzapine and clozapine result in glucose regulation abnormalities^46^. Clozapine (20 mg/kg, p.o.) nonsignificantly increased blood glucose concentrations at 30 min and 60 min, while KD025 (200 mg/kg, p.o.) resulted in a minimal change at 6 h (6 h vehicle: 50.3 ± 2.871 mg/dL; KD025 200 mg/kg: 66.2 ± 2.3 mg/dL) (Figure 5D). However, high-dose KD025 (1,000 mg/kg, p.o.) significantly increased glucose concentrations at 4 and 6 h (Figure 5D). These results suggest that at high doses, KD025 may impair glucose regulation.

**Figure 5.**
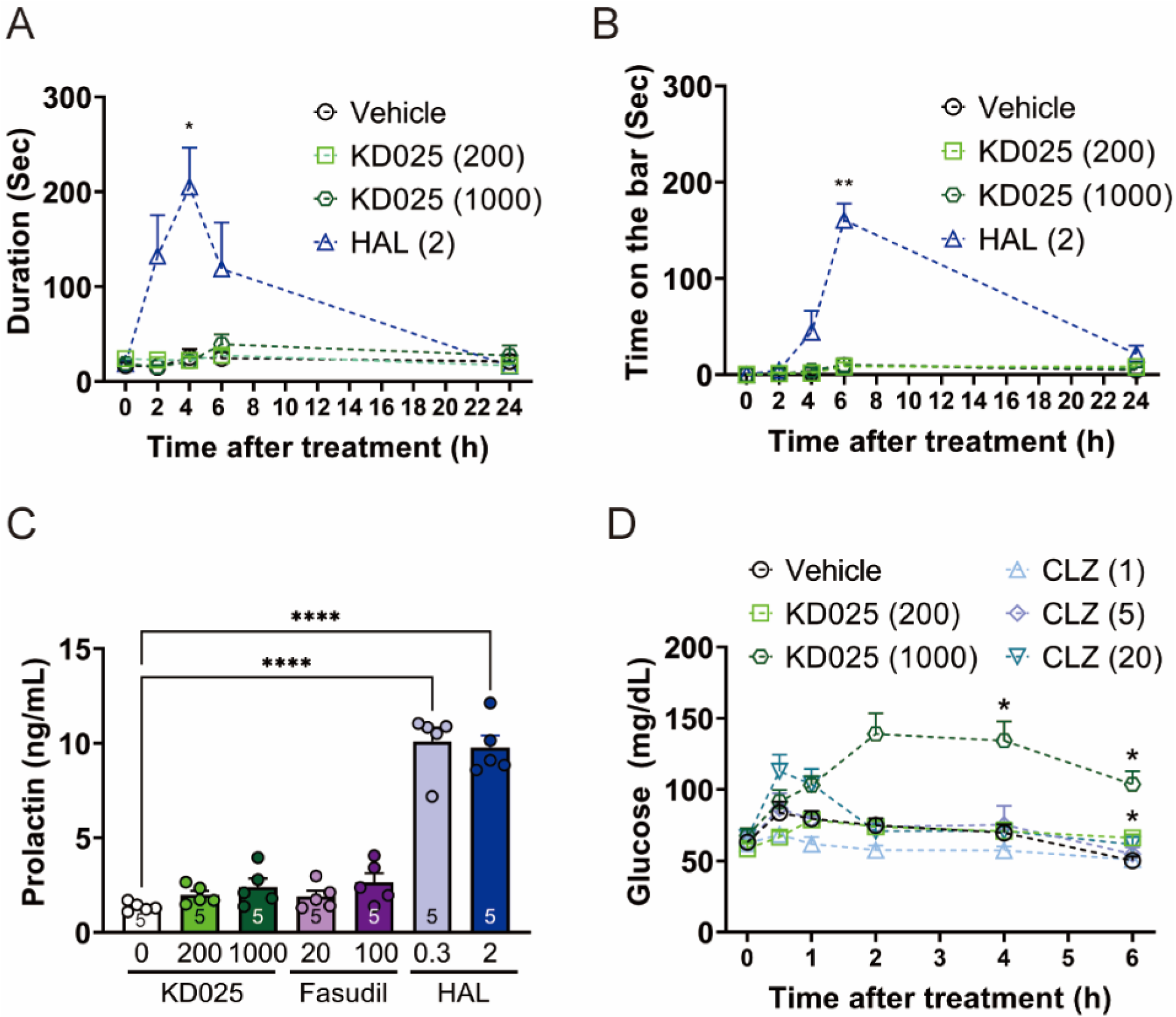
KD025 did not induce EPS or hyperprolactinemia, but did increase blood glucose levels at a high dose in naïve WT mice. A, B: The duration of T_total_ in the pole test (A) and the time on the bar in the bar test (B) before, 2, 4, 6, and 24 h after oral treatment with KD025 (200 or 1,000 mg/kg) or haloperidol (2 mg/kg). Data represent the mean + SEM (n = 4–5 WT mice per group) and were analyzed by Tukey’s multiple comparison test (*P < 0.05 and **P < 0.01 compared with the vehicle-treated group). C: Prolactin concentrations in serum 1 h after p.o. of KD025 (200 and 1,000 mg/kg), fasudil (20 and 100 mg/kg), or haloperidol (0.3 and 2 mg/kg). Data are represented as the mean + SEM (n = 5 WT mice per group) and were analyzed by Tukey’s multiple comparison test (****P < 0.0001 compared with the vehicle-treated group). D: Blood glucose concentrations before and 0.5, 1, 2, 4, and 6 h after oral treatment with KD025 (200 and 1,000 mg/kg) or clozapine (1, 5, and 20 mg/kg). Data represent the mean + SEM (n = 3–5 WT mice per group) and were analyzed by Tukey’s multiple comparison test (*P < 0.05 compared with the vehicle-treated group). HAL = haloperidol. CLZ = clozapine.

## 5 Discussion

We previously reported that the non-selective ROCK inhibitor fasudil has antipsychotic-like effects in both genetic (*Arhgap10* S490P/NHEJ mice) and pharmacological (METH- or MK-801– treated mice) models of schizophrenia^19, 21, 22^. There are two subtypes of ROCK, namely ROCK1 and ROCK2. Fasudil inhibits both subtypes with similar affinities^24^. In the present study, we aimed to optimize the use of a ROCK as a novel pharmacological target in schizophrenia, and focused on the effects of a selective ROCK2 inhibitor, KD025, in mouse models of schizophrenia. We demonstrated that ROCK2 was expressed predominantly in neurons of the mPFC and striatum. Interestingly, KD025 showed antipsychotic-like effects but only minimally induced hypotension, in contrast to the non-selective ROCK inhibitor fausdil (Figures 1–3, 4J). Moreover, KD025 exhibited minimal induction of EPS and hyperprolactinemia, both of which are typical adverse effects caused by current agents such as D2R antagonists (Figure 5).

*ROCK1* encodes 1,354 amino acids, whereas *ROCK2* encodes 1,388 amino acids; these subtypes are 64% homologous, with their kinase domains exhibiting the highest homology (92%)^23^. Thus, ROCK1 and ROCK2 show similar kinase activity in vitro^47, 48^. However, it has been reported that the expression of these subtypes varies between tissues^47^. Whereas ROCK1 mRNA is predominantly expressed in blood cells and the thymus, ROCK2 mRNA is abundantly expressed in the brain and heart of mice and humans, according to studies involving northern blotting and a human tissue–specific gene expression database, respectively^23, 25^. In the brain, ROCK2 is highly expressed in neurons, while ROCK1 is mainly distributed in glial cells^49^. By contrast, in vitro analysis showed that in both mice and humans, ROCK2 mRNA is more highly expressed in glial cells than in neurons^50^. We confirmed that ROCK2 was highly expressed in neurons, but not in astrocytes or microglia (Supplementary Figure S1). In addition, pMYPT1 was highly co-expressed with NeuN in this study (Figure 1A, 2A) and in our previous reports^19, 21^. Our observations strongly suggest that ROCK2 is highly expressed in neurons in the mouse brain. Thus, selective inhibition of ROCK2 is a promising, neuron-specific therapeutic strategy for schizophrenia.

ROCK directly phosphorylates MYPT1, at Thr696, and the myosin light chain (MLC)^51–53^. In addition, pMYPT1 (Thr696), which is the inactive, phosphorylated form, inhibits dephosphorylation of MLC, leading to sustained phosphorylation of MLC and actomyosin contractility^51–53^. In neurons, RhoA/ROCK signaling promotes spine shrinkage and destabilization via phosphorylation of MLC^54, 55^. *Arhgap10* S490P/NHEJ mice show decreased spine density in the mPFC^15^, which is consistent with the finding that schizophrenia patients exhibit a reduced density of spines in the pyramidal neurons of the prefrontal cortex^56, 57^. We showed that KD025 (p.o.), like fasudil^19^, suppressed the increased phosphorylation of MYPT1 in neurons and completely ameliorated the decreased spine density of layer 2/3 pyramidal neurons in the mPFC of *Arhgap10* S490P/NHEJ mice (Figure 1A–D). These findings are similar to those of a previous report showing that KD025 mitigated the decreased spine density in protocadherin 17–overexpressing cultured neurons^58^. These observations suggest that loss of ARHGAP10 function caused by *Arhgap10* variants results in abnormal ROCK2 activation, leading to decreased spine density of layer 2/3 pyramidal neurons in the mPFC. Importantly, KD025 diminished the decrease in spine density in adult *Arhgap10* S490P/NHEJ mice (Figure 1C, D). Thus, ROCK2 could be a therapeutic target to ameliorate the decreased spine density in the mPFC of schizophrenia patients with *ARHGAP10* gene variants.

Cognitive dysfunction is present in over 80% of schizophrenia patients, and is often observed prior to its onset^2, 59, 60^. In addition, the severity of cognitive impairment in first-episode schizophrenia is greater than that in other psychiatric disorders such as bipolar disorder and depression, and is comparable to that in patients with chronic schizophrenia^8, 60–62^. Using both genetic and pharmacological mouse models of acute schizophrenia that resemble first-episode schizophrenia^63^, we examined the effects of KD025 on cognitive dysfunction in a touchscreen-based VD task (Figures 1F, 2C, 3A). The results suggest that KD025 might target cognitive dysfunction in schizophrenia patients, regardless of the presence of *ARHGAP10* variants. The touchscreen-based VD task is a cognitive assessment method whose results translate between rodents and humans^64^. Performing the task depends on neuronal activity in the corticostriatal circuit^65^. We confirmed that KD025, like fasudil^21^, suppressed the METH-induced increase in the phosphorylation of MYPT1 in the striatum of WT mice (Figure 2A, B). This is consistent with the finding that amphetamine treatment activates RhoA in slices of mouse midbrain and in the neuroblastoma cell line SK-N-SH^66^. In addition, we previously demonstrated that fasudil suppressed the abnormal, METH-induced increase in c-Fos expression in the mPFC and striatum of *Arhgap10* S490P/NHEJ and WT mice^19–21^. Acute MK-801 treatment was also shown to induce abnormal neuronal activation in the mPFC in WT mice^67^. These observations suggest that METH- and MK-801–induced abnormal ROCK2 activation may mediate neuronal activation in the corticostriatal circuit, resulting in cognitive impairment. Taken together, KD025 might have therapeutic potential for cognitive dysfunction in patients with first-episode schizophrenia, regardless of the presence of *ARHGAP10* variants.

We also found that KD025 ameliorated the impairment of novel object recognition induced by MK-801 (Figure 3A–F). In contrast to cognitive function, KD025 had different effects on hyperlocomotion depending on whether it was caused by MK-801 or METH; KD025 suppressed hyperlocomotion resulting from MK-801, but had minimal effect on that secondary to METH (Figure 3G, H, and Supplementary Figure S2). These results are consistent with our previous report showing that fasudil suppressed hyperlocomotion induced by MK-801, but not by METH^21, 22^. In addition, KD025 failed to ameliorate the MK-801–induced impairment of PPI in WT mice (Supplementary Figure S3), while fasudil partially did so^22^. These results suggest that the activation of ROCK2 signaling plays a particularly important role in cognitive impairment in schizophrenia.

Supporting the potential clinical application of selective ROCK2 inhibitors, in naïve WT mice we confirmed that KD025 (100–300 mg/kg) had little effect on not only anxiety and general behavior but also on cognitive function, pMYPT1 level, and spine density in the mPFC (Figures 1–3, 4A–H). Of note, KD025 minimally impacted spontaneous locomotor activity (Figure 4I), although fasudil decreased it at a high dose of 30 mg/kg (i.p.)^22^. In addition, we confirmed that KD025 had little effect on systolic blood pressure in WT mice, while fasudil significantly decreased it (Figure 4J). ROCK causes vasoconstriction by promoting actomyosin contractility through MYPT1 and MLC phosphorylation in vascular smooth muscle cells and negatively regulating nitric oxide production in endothelial cells^27^. Fasudil decreases systolic blood pressure in humans and rodents^26, 68, 69^, whereas in a clinical setting, only 4.3% of KD025-treated patients with cGVHD exhibited hypotension^70^. Although ROCK2 is highly expressed in the heart as well as the brain^23, 25^, KD025 did not affect heart rate at the effective dosage (Supplementary Figure S4). Taken together, selective ROCK2 inhibitors have a smaller effect on the cardiovascular system than fasudil, while the effects on cognitive dysfunction are comparable between non-selective and selective ROCK2 inhibitors.

The adverse effects of current antipsychotics are a major problem, often leading to the discontinuation of these drugs in clinical practice^5^. EPSs, including acute dystonia, akathisia, parkinsonism, and tardive dyskinesia, are induced by D2R antagonists such as haloperidol, as well as by serotonin and dopamine antagonists such as zotepine and lurasidone^5, 71^. In addition, antipsychotics induce hyperprolactinemia by inhibiting D2R^44^. At both effective and high doses, KD025 did not induce EPS or hyperprolactinemia, while haloperidol did (Figure 5A–C, Supplemental Figure S5). Of note, haloperidol at a dose of 0.3 mg/kg partially normalized MK-801–induced hyperlocomotion and also increased the serum prolactin concentration (Figure 5C). Therefore, ROCK2 might be a more appropriate therapeutic target for schizophrenia, with fewer adverse effects than those caused by D2R antagonists. Abnormal glucose regulation is also among the adverse effects induced by SGAs such as olanzapine and clozapine^46^. SGAs block various neurotransmitter receptors, such as 5-HT2A, 5-HT2C, H1, D2, and D3, in the skeletal muscle and pancreatic β-cells and α-cells, resulting in hyperglycemia^72, 73^. We confirmed that clozapine (20 mg/kg, p.o.) increased blood glucose concentrations (Figure 5D).

By contrast, KD025 (200 mg/kg, p.o.) at an effective dose only slightly increased blood glucose concentrations, while KD025 at a high dose of 1,000 mg/kg significantly increased them in naïve WT mice (Figure 5D). In two clinical trials, the first examining KD025 and the other evaluating another ROCK2-selective inhibitor, BN101, about 10% of the population receiving KD025 exhibited hyperglycemia^70, 74^. While there are no reports in rodents confirming that ROCK2 does not induce hyperglycemia, both the non-selective ROCK inhibitor Y-27632 and ROCK1 knockout caused insulin resistance by suppressing the phosphorylation of insulin receptor substrate proteins at Ser632/635^75–77^. However, there are also reports that fasudil and Y-27632 improved glucose tolerance in a mouse model of obesity^78, 79^. In addition, there are controversial reports about the mediation of insulin secretion by ROCK^78, 80^. Of note, KD025 is known to inhibit casein kinase 2, which modulates adipocyte insulin signaling (IC50 = 50 nM)^81, 82^. Considering these observations, KD025 has minimal effects on blood glucose concentrations at its effective dose, but monitoring these concentrations might be desirable in clinical practice.

The present study has two primary limitations. The first is that we evaluated the acute effects of KD025 treatment. Schizophrenia symptoms continue throughout life, and antipsychotics are used on a chronic basis^1, 5^. A clinical trial confirmed that chronic administration of KD025 was safe for the treatment of cGVHD^28^. In the present study, 1-week treatment of KD025 recovered the abnormal spine density in the mPFC of *Arhgap10* S490P/NHEJ mice (Figure 1C, D). To develop KD025 as a novel therapeutic for schizophrenia, its antipsychotic effects and safety must be evaluated during long-term treatment in animal models of schizophrenia. The second limitation is that we were able to evaluate some of the potential adverse effects of KD025. A pilot study in animals showed that KD025 was associated with maternal toxicity and embryo-fetal developmental abnormalities^83^. Infection was reported as an adverse event seen with KD025 treatment in a clinical setting^28^. Thus, to develop selective ROCK2 inhibitors as a novel therapeutic strategy in schizophrenia, it is critical to focus not only on their beneficial effects such as antipsychotic activity and cognitive improvement, but also on adverse effects such as fetal toxicity and infection. To reduce the potential for these and other adverse effects, the neuron-specific molecules downstream of ROCK2 should be investigated.

## 6 Conclusion

In conclusion, our results suggest that targeting ROCK2 inhibition may result in new therapeutic approaches for the treatment of schizophrenia.

## Acknowledgments

Fasudil hydrochloride was kindly provided by Asahi Kasei Pharma (Tokyo, Japan). The authors wish to thank the Division for Medical Research Engineering (Eri Yorifugi and Mayumi Furukawa) at the Nagoya University Graduate School of Medicine for the use of a BZ9000 bright-field microscopic (KEYENCE), a cryostat (CM3050S; Leica), an AX-R (Nikon), and NIS-Element analysis (Nikon). The authors are also grateful to the staff of the Division of Experimental Animals at Nagoya University Graduate School of Medicine for their technical support.

## 10 Author Contributions

R.T. wrote the main text and prepared the figures, performed immunohistochemistry and Golgi staining, measured the concentration of prolactin and glucose, and analyzed the data; R.T., JZ.L., Y.L., W.Z., K.F., M.K., and M.S. performed behavioral experiments and analyzed the data; D.M. generated the *Arhgap10* S490P/NHEJ mice; R.T., A.M., and H.K. measured blood pressure and analyzed the data. JZ.L., D.T., Y.K., T. M., Ta.N., To.N., K.K., N.O., H.M., and K.Y. supervised the overall project. All authors have carefully read the paper and approved the final manuscript.

## Abbreviations

cGVHD: chronic graft-versus-host disease

CNVs: copy number variants

D2R: dopamine D2 receptor

EPS: extrapyramidal symptom

FGAs: first-generation antipsychotics

i.p.: intraperitoneal administration

GFAP: glial fibrillary acidic protein

mPFC: medial prefrontal cortex

METH: methamphetamine

MLC: myosin light chain

MYPT1: myosin phosphatase–targeting subunit 1

NeuN: neuronal nuclei

NHEJ: non-homologous end joining

NORT: novel object recognition test

pMYPT1: phosphorylated MYPT1

p.o.: oral administration

PPI: pre-pulse inhibition

ROI: region of 69 interest

ROCK: Rho-kinase

SGAs: second-generation antipsychotics

VD: visual discrimination

WT: wild-type

