## Supplemental Table S1-3 for "Antipsychotic-like effects of the selective Rho-kinase 2 inhibitor KD025 in genetic and pharmacological mouse models of schizophrenia"

Affiliations

Kiyofumi Yamada, PhD

Department of Neuropsychopharmacology and Hospital Pharmacy, Nagoya University

Address: 65 Tsurumai, Showa, Nagoya, Aichi, 466-8560, Japan

ORCID: http://orcid.org/0000-0002-5280-5180

† Rinako Tanaka and Jingzhu Liao contributed equally to this work.

### Supplementary Table S1 Summary of schedule of drug treatment

| Figure | METH (i.p.) | MK-801 (i.p.) | KD025 (p.o.) | Fasudil | Haloperidol (p.o.) | Clozapine (p.o.) |
|---|---|---|---|---|---|---|
| **1A, B** | - | - | 200 mg/kg, 120 min before | - | - | - |
| **1C, D** | - | - | 200 mg/kg/day, for 7 days | - | - | - |
| **1F, G** | 0.3 mg/kg, 30 min before | - | 50–200 mg/kg, 120 min before | - | - | - |
| **1H, I** | 0.3 mg/kg, 30 min before | - | 200 mg/kg, 120 min before | - | - | - |
| **2A, B** | 1 mg/kg, 30 min before | - | 200 mg/kg, 120 min before | - | - | - |
| **2C, D** | 1 mg/kg, 30 min before | - | 50–200 mg/kg, 120 min before | - | - | - |
| **3A, B** | - | 0.1 mg/kg, 30 min before | 50–200 mg/kg, 120 min before | - | - | - |
| **3C−F** | - | 0.1 mg/kg, 30 min before | 100–300 mg/kg, 120 min before | - | 0.3 mg/kg, 60 min before | 5 mg/kg, 60 min before |
| **3G, H** | - | 0.3 mg/kg, 0 min before | 200 mg/kg, 120 min before | - | 0.3 mg/kg, 60 min before | 5 mg/kg, 60 min before |
| **4A−H** | - | - | 100–300 mg/kg, 120 min before | - | - | - |
| **4I** | - | - | 100–300 mg/kg, 0 min before | - | - | - |
| **4J** | - | - | 100–300 mg/kg, 120 min before | 20 mg/kg, p.o. or i.p., 20 min before | - | - |
| **5A, B** | - | - | 200, 1000 mg/kg, 0 min before | - | 2 mg/kg, 0 min before | - |
| **5C** | - | - | 200, 1000 mg/kg, 1 h before | 20, 100 mg/kg, p.o., 1 h before | 0.3, 2 mg/kg, 1 h before | - |
| **5D** | - | - | 200, 1000 mg/kg, 0 min before | - | - | 1, 5, 20 mg/kg, 0 min before |
| **S2** | 2 mg/kg, 0 min before | - | 200–300 mg/kg, 120 min before | - | 0.3 mg/kg, 60 min before | 5 mg/kg, 60 min before |
| **S3** | - | 0.2 mg/kg, 30 min before | 100–300 mg/kg, 120 min before | - | 0.3 mg/kg, 60 min before | 5 mg/kg, 60 min before |
| **S4** | - | - | 100–300 mg/kg, 120 min before | 20 mg/kg, p.o. or i.p., 20 min before | - | - |
| **S5** | - | - | 200, 1000 mg/kg, 0 min before | - | 2 mg/kg, 0 min before | |

Supplementary Table S2 List of antibodies used in the previous study

| Figure | 1st antibody | Host | Catalog No. | RRID | Source | Dilutions |
|---|---|---|---|---|---|---|
| **1A, 2A** | anti–phospho-MYPT1 (Thr696) | rabbit | ABS45 | AB_10562238 | Millipore | 1:100 |
| **1A, 2A, S1A** | anti–NeuN | mouse | MAB377 | AB_2298772 | Millipore | 1:500 |
| **S1A, D** | anti–ROCK2 | rabbit | Ab125025 | AB_10972853 | Abcam | 1:200 |
| **S1D** | anti–Iba-1 | goat | NB100-1028 | AB_521594 | Novus Biologicals | 1:250 |
| **S1D** | anti–GFAP | mouse | G3893 | AB_477010 | Millipore | 1:400 |
| **Figure** | **2nd antibody** | **Host** | **Catalog No.** | **RRID** | **Source** | **Dilutions** |
| **1A, 2A** | anti–mouse Alexa Fluor 488 | goat | A11029 | AB_ 2534088 | Thermo Fisher Scientific | 1:1000 |
| **S1A** | anti–rabbit Alexa Fluor 488 | goat | A11034 | AB_2576217 | Thermo Fisher Scientific | 1:2000 |
| **S1D** | anti–rabbit Alexa Fluor 488 | donkey | A21206 | AB_2535792 | Thermo Fisher Scientific | 1:2000 |
| **S1A** | anti–mouse Alexa Fluor 568 | goat | A11031 | AB_144696 | Thermo Fisher Scientific | 1:1000 |
| **S1D** | anti–goat Alexa Fluor 568 | donkey | A11057 | AB_2534104 | Thermo Fisher Scientific | 1:1000 |
| **1A, 2A** | anti–rabbit Alexa Fluor 594 | goat | A11037 | AB_ 2534095 | Thermo Fisher Scientific | 1:1000 |
| **S1D** | anti–mouse Alexa Fluor 647 | donkey | A32787 | AB_2762830 | Thermo Fisher Scientific | 1:1000 |

### Supplementary Table S3 Summary of statistical analysis

| Figure | Number of samples | Test used | Degree of freedom and F/t/P value | Post-hoc test | Significance |
|---|---|---|---|---|---|
| **1A, B** | WT-Vehicle = 12 slices from 4 mice (male 10–13 weeks old)<br>WT-KD025 200 mg/kg = 12 slices from 4 mice (male 10–13 weeks old)<br>*Arhgap10* S490P/NHEJ-Vehicle = 12 slices from 4 mice (male 10–13 weeks old)<br>*Arhgap10* S490P/NHEJ-KD025 200 mg/kg = 12 slices from 4 mice (male 10–13 weeks old) | Two-way ANOVA | KD025: F (1, 44) = 6.013, P = 0.0182,<br>Genotype: F (1, 44) = 0.7436, P = 0.3932<br>KD025 × Genotype: F (1, 44) = 3.389, P = 0.0724 | Tukey's multiple comparisons test | Vehicle:Arhgap10 S490P/NHEJ vs. KD025 200 mg/kg:Arhgap10 S490P/NHEJ: P = 0.0202 |
| **1C, D** | WT-Vehicle = 30 dendrites from 5 mice (male 7–8 weeks old)<br>WT-KD025 200 mg/kg = 24 dendrites from 4 mice (male 7–8 weeks old)<br>*Arhgap10* S490P/NHEJ-Vehicle = 30 dendrites from 5 mice (male 7–8 weeks old)<br>*Arhgap10* S490P/NHEJ-KD025 200 mg/kg = 30 dendrites from 5 mice (male 7–8 weeks old) | Two-way ANOVA | KD025: F (1, 110) = 27.07, P < 0.0001,<br>Genotype: F (1, 110) = 5.572, P = 0.0200<br>KD025 × Genotype: F (1, 110) = 12.91, P = 0.0005 | Tukey's multiple comparisons test | Vehicle:WT vs. Vehicle:*Arhgap10* S490P/NHEJ: P = 0.0002,<br>Vehicle:*Arhgap10* S490P/NHEJ vs. KD025 200 mg/kg:*Arhgap10* S490P/NHEJ: P < 0.0001 |
| **1F** | *Arhgap10* S490P/NHEJ Saline-Vehicle = 10 (male 8~ weeks old)<br>*Arhgap10* S490P/NHEJ Saline-KD025 200 mg/kg = 10 (male 8~ weeks old)<br>*Arhgap10* S490P/NHEJ METH 0.3 mg/kg-Vehicle = 10 (male 8~ weeks old)<br>*Arhgap10* S490P/NHEJ METH 0.3 mg/kg-KD025 50 mg/kg = 10 (male 8~ weeks old)<br>*Arhgap10* S490P/NHEJ METH 0.3 mg/kg-KD025 100 mg/kg = 10 (male 8~ weeks old)<br>*Arhgap10* S490P/NHEJ METH 0.3 mg/kg-KD025 200 mg/kg = 10 (male 8~ weeks old) | One-way ANOVA | F (5, 54) = 12.06, P < 0.0001 | Tukey's multiple comparisons test | Saline-Vehicle vs. METH-Vehicle: P < 0.0001, METH-Vehicle vs. METH-KD025 100 mg/kg: P = 0.0082, METH-Vehicle vs. METH-KD025 200 mg/kg: P < 0.0001 |
| **1G** | | Kruskal-Wallis test | P = 0.9629 | N.A. | N.A. |
| **1H** | WT-Saline-Vehicle = 6 (male 8~ weeks old)<br>WT-Saline-KD025 200 mg/kg = 6 (male 8~ weeks old)<br>WT-METH 0.3 mg/kg-Vehicle = 6 (male 8~ weeks old)<br>WT-METH 0.3 mg/kg-KD025 200 mg/kg = 6 (male 8~ weeks old) | Two-way ANOVA | METH: F (1, 5) = 1.134, P=0.3355,<br>KD025: F (1, 5) = 2.738, P=0.1589<br>METH × KD025: F (1, 5) = 0.02262, P=0.8863 | N.A. | N.A. |
| **1I** | | Kruskal-Wallis test | P = 0.0743 | N.A. | N.A. |
| **2AB** | WT-Saline-Vehicle = 18 slices from 6 mice (male 12 weeks old)<br>WT-Saline-KD025 200 mg/kg = 18 slices from 6 mice (male 12 weeks old)<br>WT-METH 1 mg/kg-Vehicle = 18 slices from 6 mice (male 12 weeks old)<br>WT-METH 1 mg/kg-KD025 200 mg/kg = 18 slices from 6 mice (male 12 weeks old) | Two-way ANOVA | METH: F (1, 68) = 34.96, P < 0.0001,<br>KD025: F (1, 68) = 13.84, P = 0.0004,<br>METH × KD025: F (1, 68) = 0.2088, P = 0.6492 | Tukey's multiple comparisons test | Saline:Vehicle vs. METH:Vehicle: P = 0.0002,<br>METH-Vehicle vs. METH:KD025 200 mg/kg: P = 0.0219 |
| **2C** | WT-Saline-Vehicle = 10 (male 8~ weeks old)<br>WT-Saline-KD025 200 mg/kg = 9 (male 8~ weeks old)<br>WT-METH 1 mg/kg-Vehicle = 10 (male 8~ weeks old)<br>WT-METH 1 mg/kg-KD025 50 mg/kg = 9 (male 8~ weeks old)<br>WT-METH 1 mg/kg-KD025 100 mg/kg = 9 (male 8~ weeks old)<br>WT-METH 1 mg/kg-KD025 200 mg/kg = 10 (male 8~ weeks old) | One-way ANOVA | F (5, 51) = 5.160, P = 0.0007 | Tukey's multiple comparisons test | Saline-Vehicle vs. METH-Vehicle: P = 0.0023, METH-Vehicle vs. METH-KD025 200 mg/kg: P = 0.0098 |
| **2D** | | Kruskal-Wallis test | P = 0.0391 | Dunn's multiple comparisons test t | N.S. |
| **3A** | WT-Saline-Vehicle = 10 (male 8~ weeks old)<br>WT-Saline-KD025 200 mg/kg = 7 (male 8~ weeks old)<br>WT-MK-801 0.1 mg/kg-Vehicle = 8 (male 8~ weeks old)<br>WT-MK-801 0.1 mg/kg-KD025 50 mg/kg = 8 (male 8~ weeks old)<br>WT-MK-801 0.1 mg/kg-KD025 100 mg/kg = 6 (male 8~ weeks old)<br>WT-MK-801 0.1 mg/kg-KD025 200 mg/kg = 9 (male 8~ weeks old) | One-way ANOVA | F (5, 42) = 6.963, P < 0.0001 | Tukey's multiple comparisons test | Saline-Vehicle vs. MK-801-Vehicle: P = 0.0008, MK-801-Vehicle vs. MK-801-KD025 100 mg/kg: P = 0.0177, MK-801-Vehicle vs. MK-801-KD025 200 mg/kg: P < 0.0001 |
| **3B** | | Kruskal-Wallis test | P = 0.6721 | N.A. | N.A. |
| **3C** | WT-Saline-Vehicle = 16 (male 10–12 weeks old)<br>WT-Saline-KD025 300 mg/kg = 10 (male 10–12 weeks old)<br>WT-MK-801 0.1 mg/kg-Vehicle = 16 (male 10–12 weeks old)<br>WT-MK-801 0.1 mg/kg-KD025 100 mg/kg = 12 (male 10–12 weeks old)<br>WT-MK-801 0.1 mg/kg-KD025 200 mg/kg = 13 (male 10–12 weeks old)<br>WT-MK-801 0.1 mg/kg-KD025 300 mg/kg = 10 (male 10–12 weeks old)<br>WT-MK-801 0.1 mg/kg-Haloperidol 0.3 mg/kg = 10 (male 10–12 weeks old)<br>WT-MK-801 0.1 mg/kg-Clozapine 5 mg/kg = 10 (male 10–12 weeks old) | One-way ANOVA | F (7, 89) = 1.027, P = 0.4182 | N.A. | N.A. |
| **3D** | | One-way ANOVA | F (7, 89) = 18.75, P < 0.0001 | Tukey's multiple comparisons test | Saline-Vehicle vs. MK-801-Vehicle: P < 0.0001, MK-801-Vehicle vs. MK-801-KD025 200 mg/kg: P < 0.0001, MK-801-Vehicle vs. MK-801-KD025 300 mg/kg: P < 0.0001, MK-801-Vehicle vs. MK-801-CLZ 5 mg/kg: P < 0.0001 |
| **3E** | | One-way ANOVA | F (7, 89) = 4.297, P = 0.0004 | Tukey's multiple comparisons test | MK-801-Vehicle vs. MK-801-HAL 0.3 mg/kg: P < 0.0001 |
| **3F** | | One-way ANOVA | F (7, 89) = 0.9835, P = 0.4484 | N.A. | N.A. |
| **3G** | WT-Saline-Vehicle = 10 (male 11–12 weeks old)<br>WT-MK-801 0.3 mg/kg-Vehicle = 10 (male 11–12 weeks old)<br>WT-MK-801 0.3 mg/kg-KD025 200 mg/kg = 10 (male 11–12 weeks old)<br>WT-MK-801 0.3 mg/kg-Haloperidol 0.3 mg/kg = 10 (male 11–12 weeks old)<br>WT-MK-801 0.3 mg/kg-Clozapine 5 mg/kg = 10 (male 11–12 weeks old) | Two-way Repeated measures (RM) ANOVA | Time: F (23, 1035) = 45.15, P < 0.0001,<br>Drugs: F (4, 45) = 20.69, P < 0.0001,<br>Time × Drugs: F (92, 1035) = 4.376, P < 0.0001 | Tukey's multiple comparisons test | Saline-Vehicle vs. MK-801-Vehicle: P < 0.05 at time points of 5–95 min, MK-801-Vehicle vs. MK-801-KD025 200 mg/kg: P < 0.05 at time points of 10, 30, 40–55 min, MK-801-Vehicle vs. MK-801-HAL 0.3 mg/kg: P < 0.05 at time points of 5–20, 40–90 min |
| **3H** | | One-way ANOVA | F (4, 45) = 20.69, P < 0.0001 | Tukey's multiple comparisons test | Saline-Vehicle vs. MK-801-Vehicle: P < 0.0001, MK-801-Vehicle vs. MK-801-KD025 200 mg/kg: P = 0.0226, MK-801-Vehicle vs. MK-801-HAL 0.3 mg/kg: P < 0.0001 |

| | | | | | |
|---|---|---|---|---|---|
| **4A** | WT-Vehicle = 10 (male 8 weeks old)<br>WT-KD025 100 mg/kg = 9 (male 8 weeks old)<br>WT-KD025 200 mg/kg = 10 (male 8 weeks old)<br>WT-KD025 300 mg/kg = 9 (male 8 weeks old) | Two-way ANOVA | Zone: F (1, 68) = 8522 P < 0.0001,<br>KD025: F (3, 68) = 0.008934, P = 0.9988,<br>Zone × KD025: F (3, 68) = 1.125, P = 0.3453 | Dunnett's multiple comparisons test | N.S. |
| **4B** | | One-way ANOVA | F (3, 34) = 2.954, P = 0.0463 | Dunnett's multiple comparisons test | N.S. |
| **4C** | | One-way ANOVA | F (3, 34) = 1.282, P = 0.2963 | N.A. | N.A. |
| **4D** | | Kruskal-Wallis test | P = 0.9941 | N.A. | N.A. |
| **4E** | | Kruskal-Wallis test | P = 0.6448 | N.A. | N.A. |
| **4F** | | Kruskal-Wallis test | P = 0.7287 | N.A. | N.A. |
| **4G** | | Kruskal-Wallis test | P = 0.5424 | N.A. | N.A. |
| **4H** | | Kruskal-Wallis test | P = 0.6041 | N.A. | N.A. |
| **4I** | WT-Vehicle = 10 (male 9–10 weeks old)<br>WT-KD025 100 mg/kg = 9 (male 9–10 weeks old)<br>WT-KD025 200 mg/kg = 10 (male 9–10 weeks old)<br>WT-KD025 300 mg/kg = 10 (male 9–10 weeks old) | Two-way RM ANOVA | Time: F (2, 70) = 163.3, P < 0.0001,<br>KD025: F (3, 35) = 1.714, P = 0.1820, Time × KD025: F (6, 70) = 0.9241, P = 0.4832 | Dunnett's multiple comparisons test | N.S. |
| **4J** | WT-Vehicle = 5 (male 8–9 weeks old)<br>WT-KD025 100 mg/kg = 5 (male 8–9 weeks old)<br>WT-KD025 200 mg/kg = 4 (male 8–9 weeks old)<br>WT-KD025 300 mg/kg = 5 (male 8–9 weeks old)<br>WT-Fasudil 20 mg/kg, p.o. = 5 (male 8–9 weeks old)<br>WT-Fasudil 20 mg/kg, i.p. = 4 (male 8–9 weeks old) | Two-way RM ANOVA | pre/post: F (1, 22) = 51.71, P < 0.0001, ROCK inhibitors: F (5, 22) = 9.692, P < 0.0001, pre/post × ROCK inhibitors: F (5, 22) = 6.875, P = 0.0005 | Šídák's multiple comparisons test | Pre vs Post: Fasudil 20 mg/kg p.o.: P = 0.0001, Fasudil 20 mg/kg i.p.: P < 0.0001 |
| **5A** | WT-Vehicle = 5 (male 10 weeks old)<br>WT-KD025 200 mg/kg = 4 (male 10 weeks old)<br>WT-KD025 1000 mg/kg = 5 (male 10 weeks old)<br>WT-Haloperidol 2 mg/kg =5 (male 10 weeks old) | Two-way RM ANOVA | Time: F (2.297, 32.16) = 6.951, P = 0.0021,<br>Drug: F (3, 14) = 7.636, P = 0.0029,<br>Time ×Drug: F (12, 56) = 6.808, P < 0.0001 | Tukey's multiple comparisons test | 4 h: Vehicle vs. Haloperidol 2 mg/kg: P = 0.0380 |
| **5B** | WT-Vehicle = 5 (male 10 weeks old)<br>WT-KD025 200 mg/kg = 5 (male 10 weeks old)<br>WT-KD025 1000 mg/kg = 5 (male 10 weeks old)<br>WT-Haloperidol 2 mg/kg =5 (male 10 weeks old) | Two-way RM ANOVA | Time: F (2.008, 32.14) = 39.83, P < 0.0001,<br>Drug: F (3, 16) = 21.66, P < 0.0001,<br>Time ×Drug: F (12, 64) = 27.16, P < 0.0001 | Tukey's multiple comparisons test | 6 h: Vehicle vs. Haloperidol 2 mg/kg: P = 0.0014 |
| **5C** | WT-Vehicle = 5 (male 10 weeks old)<br>WT-KD025 200 mg/kg = 5 (male 10 weeks old)<br>WT-KD025 1000 mg/kg = 5 (male 10 weeks old)<br>WT-Fasudil 20 mg/kg. = 5 (male 10 weeks old)<br>WT-Fasudil 100 mg/kg= 5 (male 10 weeks old)<br>WT-Haloperidol 0.3 mg/kg =5 (male 10 weeks old)<br>WT-Haloperidol 2 mg/kg =5 (male 10 weeks old) | One-way ANOVA | F (6, 28) = 67.51, P < 0.0001 | Tukey's multiple comparisons test | Vehicle vs. Haloperidol 0.3 mg/kg: P < 0.0001<br>Vehicle vs. Haloperidol 2 mg/kg: P < 0.0001 |
| **5D** | WT-Vehicle = 5 (male 10 weeks old)<br>WT-KD025 200 mg/kg = 5 (male 10 weeks old)<br>WT-KD025 1000 mg/kg = 5 (male 10 weeks old)<br>WT-Clozapine 1 mg/kg =3 (male 10 weeks old)<br>WT-Clozapine 5 mg/kg =3 (male 10 weeks old)<br>WT-Clozapine 20 mg/kg =5 (male 10 weeks old) | Two-way RM ANOVA | Time: F (3.986, 79.73) = 14.29,<br>Drug: F (5, 20) = 14.02,<br>Time ×Drug: F (25, 100) = 5.573 | Tukey's multiple comparisons test | 4 h: Vehicle vs. KD025 1000 mg/kg: P = 0.0382<br>6 h: Vehicle vs. KD025 200 mg/kg: P = 0.0228<br>6 h: Vehicle vs. KD025 1000 mg/kg: P = 0.0213 |
| **S1** | WT =3 (male 16–18 weeks old) | N.A. | N.A. | N.A. | N.A. |
| **S2A** | WT-Saline-Vehicle = 10 (male 10 weeks old)<br>WT-METH 2 mg/kg-Vehicle = 10 (male 10 weeks old)<br>WT-METH 2 mg/kg-KD025 200 mg/kg = 10 (male 10 weeks old)<br>WT-METH 2 mg/kg-KD025 300 mg/kg = 13 (male 10 weeks old)<br>WT-METH 2 mg/kg -Haloperidol 0.3 mg/kg = 6 (male 10 weeks old)<br>WT-METH 2 mg/kg -Clozapine 5 mg/kg = 6 (male 10 weeks old) | Two-way RM ANOVA | Time: F (3.886, 190.4) = 25.45, P < 0.0001,<br>Drugs: F (5, 49) = 14.52, P < 0.0001,<br>Time × Drugs: F (120, 1176) = 2.625, P < 0.0001 | Tukey's multiple comparisons test | Saline-Vehicle vs. METH-Vehicle: P < 0.05 at time points of 15–105 min, METH-Vehicle vs. METH-KD025 200 mg/kg: P = 0.0384 at time points of 55 min, METH-Vehicle vs. METH-HAL 0.3 mg/kg: P < 0.05 at time points of 5–105 min |
| **S2B** | | One-way ANOVA | F (5, 49) = 14.52, P < 0.0001 | Tukey's multiple comparisons test | Saline-Vehicle vs. METH-Vehicle: P < 0.0001, METH-Vehicle vs. METH-Haloperidol 0.3 mg/kg: P < 0.0001 |
| **S3A** | WT-Saline-Vehicle = 15 (male 10–12 weeks old)<br>WT-Saline-KD025 300 mg/kg = 10 (male 10–12 weeks old)<br>WT-MK-801 0.2 mg/kg-Vehicle = 16 (male 10–12 weeks old)<br>WT-MK-801 0.2 mg/kg-KD025 100 mg/kg = 11 (male 10–12 weeks old)<br>WT-MK-801 0.2 mg/kg-KD025 200 mg/kg = 11 (male 10–12 weeks old)<br>WT-MK-801 0.2 mg/kg-KD025 300 mg/kg = 10 (male 10–12 weeks old)<br>WT-MK-801 0.2 mg/kg-Haloperidol 0.3 mg/kg = 10 (male 10–12 weeks old)<br>WT-MK-801 0.2 mg/kg-Clozapine 5 mg/kg = 10 (male 10–12 weeks old) | Two-way RM ANOVA | Prepulse: F (3, 255) = 132.6, P < 0.0001,<br>Drugs: F (7, 85) = 11.23, P < 0.0001,<br>Prepulse × Drugs: F (21, 255) = 1.254, P = 0.2076 | Tukey's multiple comparisons test | 69, 73, 77, and 81 dB: Saline-Vehicle vs. MK-801-Vehicle: P < 0.0001 |
| **S3B** | | One-way ANOVA | F (7, 85) = 1.517, P = 0.1725 | N.A. | N.A. |

| | | | | | |
|---|---|---|---|---|---|
| **S4** | WT-Vehicle = 5 (male 8–9 weeks old)<br>WT-KD025 100 mg/kg = 5 (male 8–9 weeks old)<br>WT-KD025 200 mg/kg = 4 (male 8–9 weeks old)<br>WT-KD025 300 mg/kg = 5 (male 8–9 weeks old)<br>WT-Fasudil 20 mg/kg, p.o. = 5 (male 8–9 weeks old)<br>WT-Fasudil 20 mg/kg, i.p. = 4 (male 8–9 weeks old) | Two-way RM ANOVA | pre/post: F (1, 22) = 5.149 P = 0.0334, ROCK inhibitors: F (5, 22) = 0.7178, P = 0.6169, pre/post × ROCK inhibitors: F (5, 22) = 2.850, P = 0.0393 | Šídák's multiple comparisons test | Pre vs. Post: Fasudil 20 mg/kg p.o.: P = 0.0340 |
| **S5** | WT-Vehicle = 5 (male 10 weeks old)<br>WT-KD025 200 mg/kg = 4 (male 10 weeks old)<br>WT-KD025 1000 mg/kg = 5 (male 10 weeks old)<br>WT-Haloperidol 2 mg/kg =5 (male 10 weeks old) | Two-way ANOVA | Time: F (1.518, 21.25) = 3.463, P = 0.0611,<br>Drug: F (3, 14) = 3.587, P = 0.0412,<br>Time ×Drug: F (12, 56) = 4.193, P = 0.0001 | Tukey's multiple | N.S. |
