## Supplemental figure S1-S5 for "Antipsychotic-like effects of the selective Rho-kinase 2 inhibitor KD025 in genetic and pharmacological mouse models of schizophrenia"

Affiliations

Kiyofumi Yamada, PhD

Department of Neuropsychopharmacology and Hospital Pharmacy, Nagoya University

Address: 65 Tsurumai, Showa, Nagoya, Aichi, 466-8560, Japan

ORCID: http://orcid.org/0000-0002-5280-5180

† Rinako Tanaka and Jingzhu Liao contributed equally to this work.

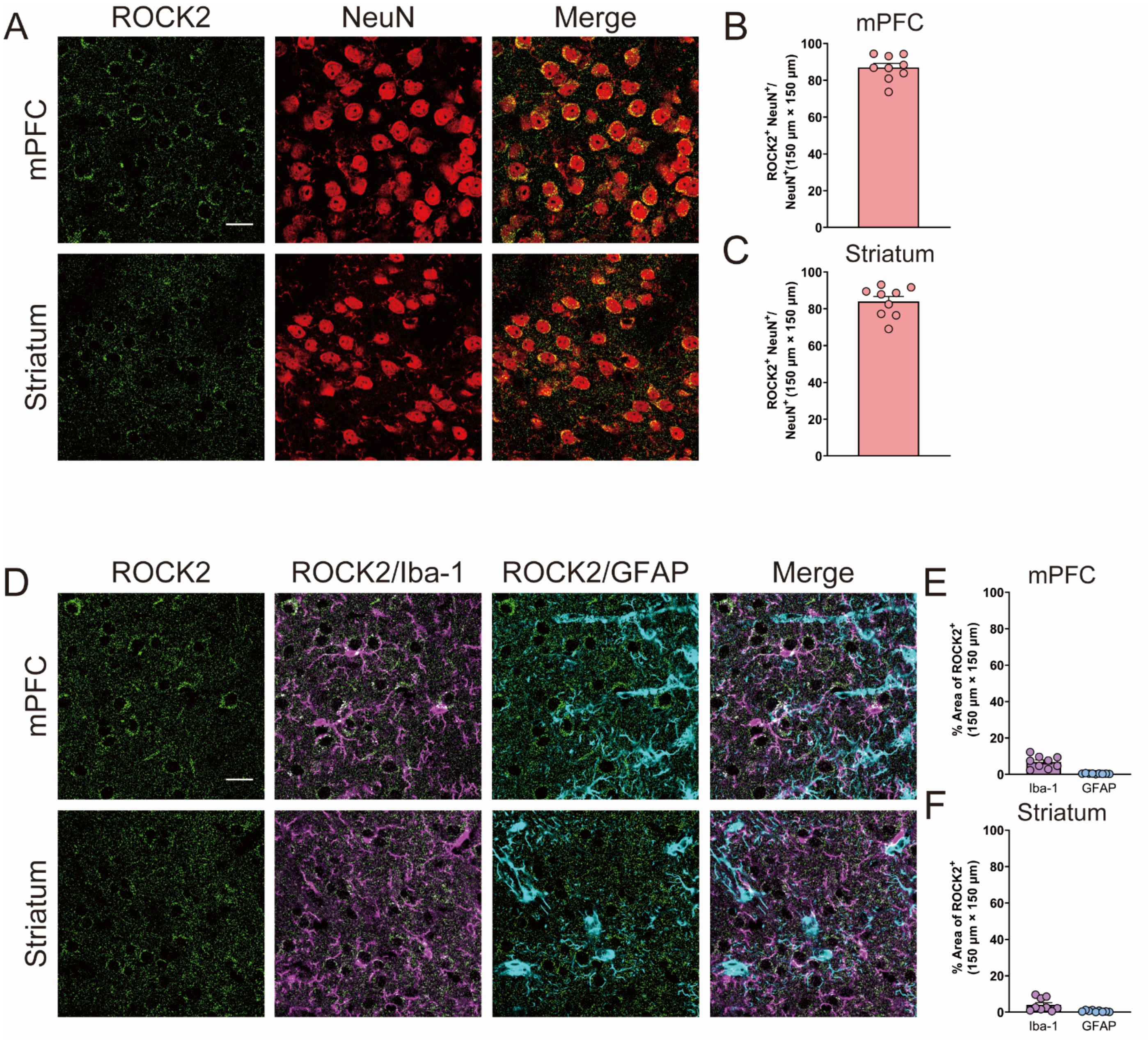


**Supplementary Figure S1 ROCK2 was expressed in neurons, but only minimally in microglia and astrocytes in the mPFC and striatum of WT mice.**

A: Representative immunoreactivity images of ROCK2 (green) and NeuN (red) in the mPFC and striatum of WT mice (scale bar indicates 20 μm). B, C: Ratio of ROCK2 and NeuN double-positive neurons to NeuN-positive neurons in the mPFC (B) and striatum (C) of WT mice. Data represent the mean + SEM (n = 9 (three mice per group, three slices per mouse)). D: Representative immunoreactivity images of ROCK2 (green), merging of ROCK2 (green) and Iba-1 (magenta), and merging of ROCK2 (green) and GFAP (cyan) in the mPFC and striatum of WT mice (scale bar indicates 20 μm). E, F: Percentage of the Iba-1- or GFAP-positive area that is double positive for ROCK2 in the mPFC (E) and striatum (F) of WT mice. Data represent the mean + SEM (n = 9 slices (three mice per group, three slices per mouse)).

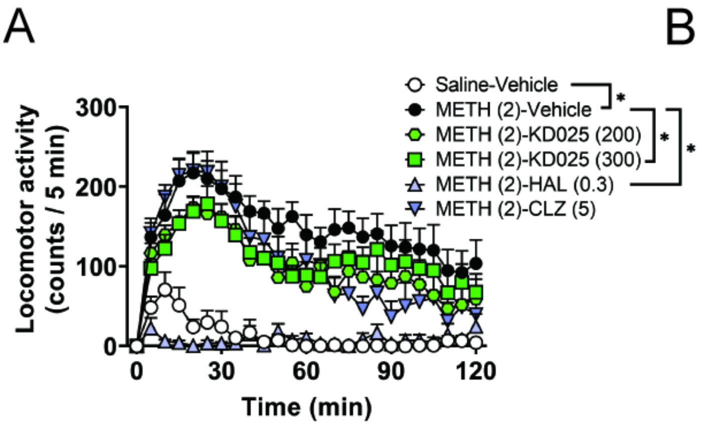


**Supplementary Figure S2 KD025 had little effect on METH-induced hyperlocomotion in WT mice.** Locomotor activity time course (A) and total locomotor activity (B) after METH treatment. Haloperidol (0.3 mg/kg, p.o.), clozapine (5 mg/kg, p.o.), or KD025 (200 or 300 mg/kg, p.o.) was administered 60, 60, or 120 min, respectively, before the test, and the locomotor activity was measured for 120 min immediately after METH treatment (2 mg/kg, i.p.). Data are represented as the mean + SEM (n = 6–13 WT mice per group) and were analyzed by Tukey's multiple comparison test ($^{*}P < 0.05$ and $^{****}P < 0.0001$). HAL = haloperidol. CLZ = clozapine.

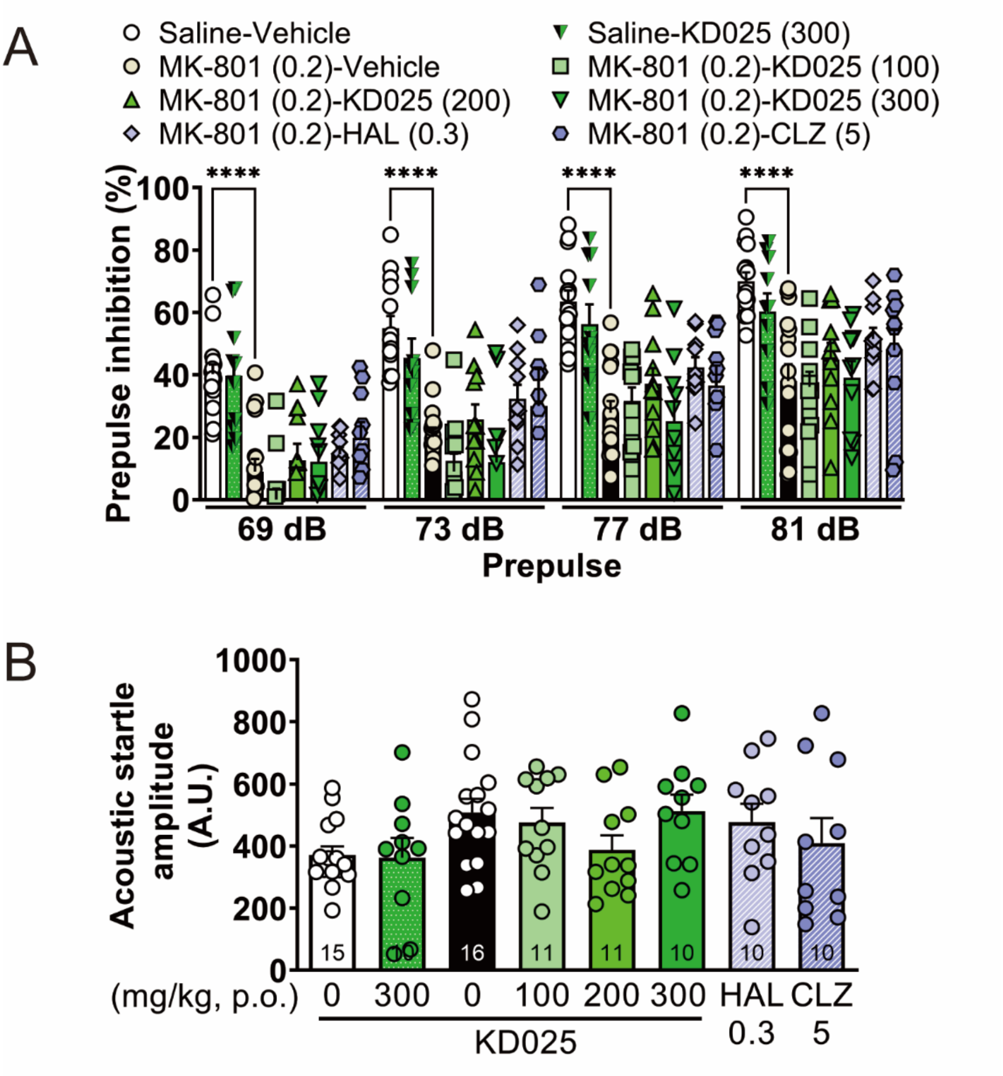


**Supplementary Figure S3 KD025 had little effect on MK-801–induced pre-pulse inhibition deficits of startle response in mice** A: Pre-pulse inhibition (%) at four different pre-pulse intensities (69, 73, 77, and 81 dB). B: Acoustic startle response measured in trials without pre-pulse. MK-801 (0.2 mg/kg, i.p.), haloperidol (0.3 mg/kg, p.o.), clozapine (5 mg/kg, p.o.), or KD025 (200 or 300 mg/kg, p.o.) was administered 30, 60, 60, or 120 min, respectively, before the test. Data are represented as the mean + SEM (n = 10–16 WT mice per group) and were analyzed by Tukey's multiple comparison test ($^{****}P < 0.0001$). HAL = haloperidol. CLZ = clozapine.

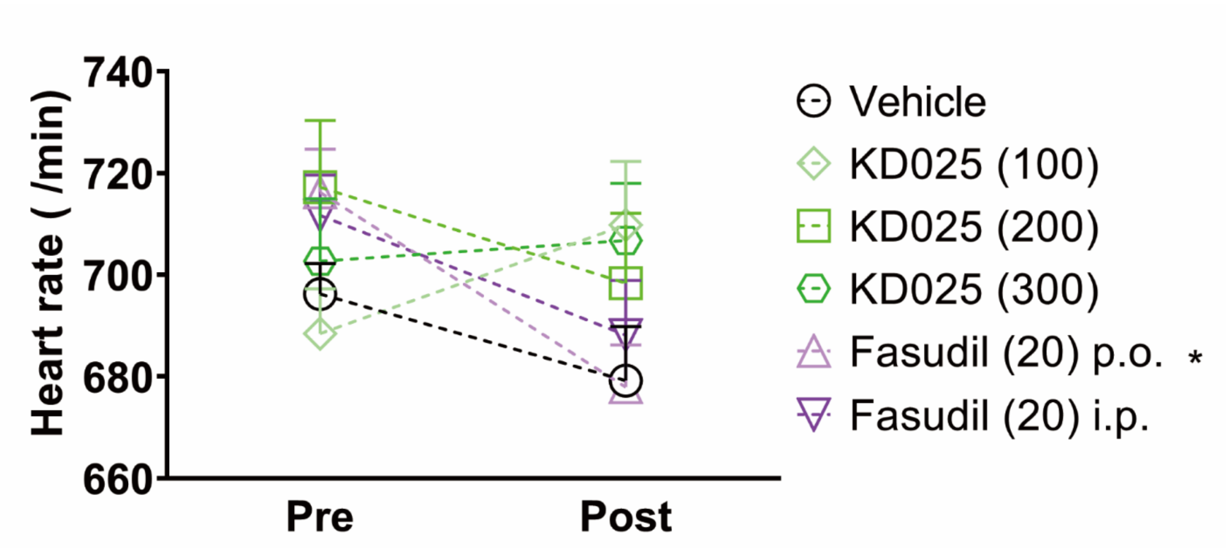


**Supplementary Figure S4 KD025 had little effect on heart rate in WT mice.**
Heart rate before fasudil (20 mg/kg, i.p. or p.o.) or KD025 (100–300 mg/kg, p.o.) treatment, and 20 and 120 min afterward, respectively. Data represent the mean + SEM (n = 4–5 WT mice per group) and were analyzed by Šídák's multiple comparisons test (*P < 0.05 compared with pre-treatment).

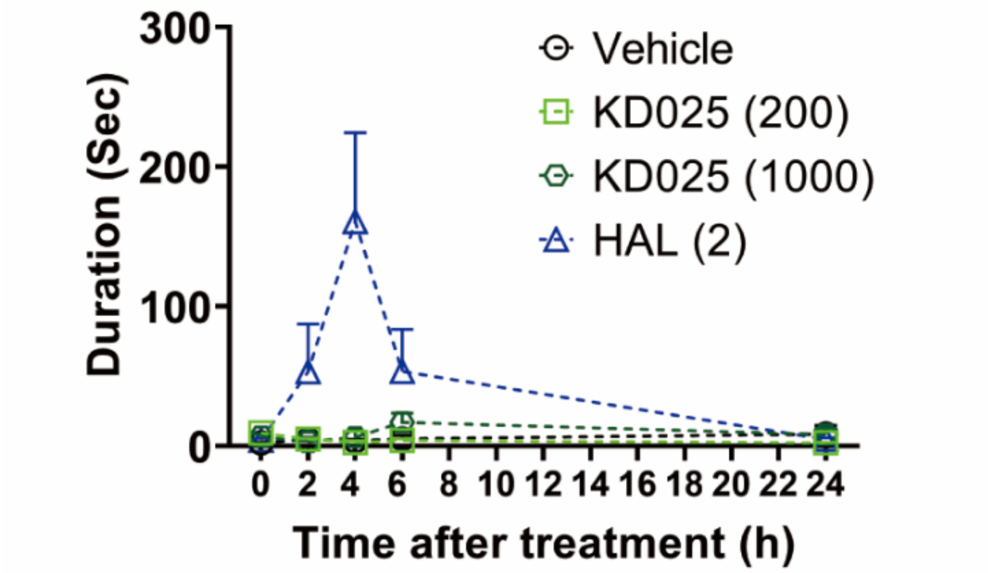


**Supplementary Figure S5 KD025 had little effect on the $T_{turn}$ duration in the pole test.**
The $T_{turn}$ duration in the pole test before and 2, 4, 6, and 24 h after oral treatment with KD025 (200 or 1,000 mg/kg) or haloperidol (2 mg/kg). Data represent the mean + SEM (n = 4–5 WT mice per group) and were analyzed by Tukey's multiple comparison test. HAL = haloperidol.
