## Supplemental material & methods for "Antipsychotic-like effects of the selective Rho-kinase 2 inhibitor KD025 in genetic and pharmacological mouse models of schizophrenia"

Affiliations

Kiyofumi Yamada, PhD

Department of Neuropsychopharmacology and Hospital Pharmacy, Nagoya University
Address: 65 Tsurumai, Showa, Nagoya, Aichi, 466-8560, Japan

ORCID: http://orcid.org/0000-0002-5280-5180

† Rinako Tanaka and Jingzhu Liao contributed equally to this work.

**Supplemental material & methods**

1. Animals

*Arhgap10* S490P/NHEJ variants mice (NHEJ and S490P line) were generated on a C57BL/6J genetic background as described previously[1]. *Arhgap10* S490P/NHEJ (n = 28) mice and their WT littermates (n = 23) were obtained by breeding 2 lines of heterozygous *Arhgap10* mutant mice (NHEJ line and S490P line). Male C57BL/6J mice (n = 432) were obtained from the Japan SLC (Shizuoka, Japan). Male mice aged over 7 weeks were used in the experiment. Mice were housed at a density of 4–6 mice per cage (28 cm length × 17 cm width × 13 cm height) in standard conditions (23 ± 1°C, 50 ± 5% humidity) with a 12-h light/dark cycle. Food and water were available ad libitum. Animals were handled in accordance with the guidelines established by the Institutional Animal Care and Use Committee of Nagoya University, the Guiding Principles for the Care and Use of Laboratory Animals approved by the Japanese Pharmacological Society, and Guide for the Care and Use of Laboratory Animals by the National Institutes of Health of the United States.

2. Drug treatment

KD025 (Slx-2119) (Cat#HY-15307/CS-0776; MedChemExpress, Monmouth Junction, NJ, USA), haloperidol (Mitsubishi Tanabe Pharma Corporation, Osaka, Japan) and clozapine (Tokyo Chemical Industry, Tokyo, Japan) were suspended in 0.1% carboxymethylcellulose–saline. Fasudil monohydrochloride salt (purity > 99%) was kindly supplied by Asahi Kasei Pharma (Tokyo, Japan) and suspended in saline. We set up the dose and timing of drug administration based on previous research[2-5]. In a pharmacokinetic analysis, KD025 was detected in the plasma and brain until 9 h after oral administration (100, 200 mg/kg)[2]. In a non-compartment analysis, the $T_{max}$ of KD025 in plasma and brain was 2 h[2]. Thus, we measured the phosphorylation levels of MYPT1 in the mPFC of *Arhgap10* S490P/NHEJ mice 2 h after oral administration of KD025 (200 mg/kg). To evaluate the phosphorylation levels of MYPT1, KD025 (200 mg/kg) or vehicle was administered orally 120 min before sample collection, and METH (Sumitomo Dainippon Pharma, Osaka, Japan) (1 mg/kg) was administered intraperitoneally 30 min before sample collection. For Golgi staining, mice were orally treated with KD025 (200 mg/kg) or vehicle once a day for 7 days in order to detect the effect on spine density, because spine turnover was reported to occur every 2 days in adult mice[6, 7]. On the day after the last KD025 administration, brain samples were collected for Golgi staining. For the behavioral experiments, KD025 (50–300 mg/kg) or vehicle was administered orally 120 min before the tests, except for the spontaneous locomotor test. Haloperidol (0.3 mg/kg), clozapine (5 mg/kg), or vehicle was administered orally 60 min before the tests. METH (0.3 and 1 mg/kg), MK-801 (0.1–0.3 mg/kg), or vehicle was administered intraperitoneally 30 min before the touchscreen-based VD task, the NORT and the prepulse inhibition test, or immediately before the hyperlocomotion test. For the tail-cuff evaluations, KD025 (100–300 mg/kg, p.o.), fasudil (20 mg/kg, p.o. or i.p.), or vehicle (p.o.) was administrated 120 min and 20 min before test. For the measurement of extrapyramidal symptoms and plasma prolactin concentrations, KD025 (200 or 1000 mg/kg), haloperidol (0.3 and 2 mg/kg), fasudil (20 and 100 mg/kg), or vehicle was administered orally. For the measurement of blood glucose concentrations, KD025 (200 or 1000 mg/kg), clozapine (1, 5, and 20 mg/kg), or vehicle was administered orally. Detailed information concerning the schedule of drug treatment is shown in Supplementary Table S1.

3. Immunohistochemistry

Immunohistochemistry was conducted as previously described[4]. In the immunohistochemical analysis of pMYPT1, mice were perfused intracardially with ice-cold 4% paraformaldehyde (PFA) in 0.1 M phosphate buffer (PB) (pH 7.4) at a rate of 10 mL/min for 6 min under anesthesia with medetomidine, midazolam, and butorphanol. In the immunohistochemical analysis of ROCK2, mice were perfused intracardially with ice-cold 0.1 M PB followed by 4 % PFA in 0.1 M PB under anesthesia with isoflurane. The brains were post-fixed with the same fixative and cryoprotected with 20% sucrose followed by 30% sucrose in 0.1 M PB. Frozen 20-µm sections for the immunohistochemical analysis were cut coronally using a cryostat (CM3050S; Leica, Wetzlar, Germany). Cryosections were fixed with 4% PFA in 0.1 M PB for 5 min and permeabilized with 0.3% Triton X-100/ phosphate-buffered saline (PBS) for 10 min in the immunohistochemical analysis of pMYPT1 or HistVTone (Cat#06380-76; Nacalai Tesque, Kyoto, Japan) for 20 min at 70°C in the immunohistochemical analysis of ROCK2. After incubation in blocking solution (5% goat or donkey serum / PBS with 0.3% Triton X-100) for 60 min, the sections were immunostained overnight with the primary antibody. The primary antibodies used were as follows: rabbit anti–phospho-

MYPT1 (Thr696) (Cat# ABS45, RRID:AB_10562238, 1:100 dilution; Millipore) and mouse anti–neuronal nuclei (NeuN) (Cat# MAB377, RRID:AB_2298772, 1:500 dilution; Millipore), anti-ROCK2 (Cat#Ab125025, RRID:AB_10972853, 1:200 dilution; Abcam, Cambridge, UK), anti–Iba-1 (Cat# NB100-1028, RRID:AB_521594, 1:250 dilution; Novus Biologicals, Centennial, USA), anti-GFAP)(Cat# G3893, AB_477010, 1:400 dilution; Millipore). After washing in PBS, the sections were incubated with the secondary antibody at room temperature for 1 h. The secondary antibodies were goat anti–mouse Alexa Fluor 488 (Cat# A11029, RRID:AB_ 2534088, 1:1000 dilution; Thermo Fisher Scientific, Waltham, MA, USA), goat anti–rabbit Alexa Fluor 488 (Cat# A11034, RRID:AB_2576217, 1:2000 dilution; Thermo Fisher Scientific, Waltham, MA, USA), donkey anti–rabbit Alexa Fluor 488 (Cat# A21206, RRID:AB_2535792, 1:2000 dilution; Thermo Fisher Scientific), goat anti–mouse Alexa Fluor 568 (Cat# A11031, RRID:AB_144696, 1:1000 dilution; Thermo Fisher Scientific), donkey anti–goat Alexa Fluor 568 (Cat# A11057, RRID:AB_2534104, 1:1000 dilution; Thermo Fisher Scientific), goat anti–rabbit Alexa Fluor 594 (Cat# A11037, 1:1000 dilution, RRID:AB_ 2534095; Thermo Fisher Scientific), donkey anti–mouse Alexa Fluor 647 (Cat# A32787, RRID:AB_2762830, 1:1000 dilution; Thermo Fisher Scientific). After washing in PBS, the sections were mounted on an adhesive silane (MAS)-coated glass slide (Matsunami, Osaka, Japan) with Fluorescent Mounting Medium (Dako, Santa Clara, CA, USA) and a coverslip. Detailed information on the antibodies used in this study is shown in Supplemental Table S2. For the immunohistochemical analysis of pMYPT1, fluorescence images were captured using a confocal laser microscope (LSM710; Carl Zeiss AG, Oberkochen, Germany) with 20×/0.8 NA objective lens, and the number of positive cells was blindly counted within each 93 μm × 93 μm ROI, using Fiji/ImageJ software package[8]. For the immunohistochemical analysis of ROCK2, fluorescence images were captured using a confocal laser microscope (AX R; Nikon, Tokyo, Japan) with a 60×/1.20 NA water-immersion objective lens, and the number of positive cells and the signal area were measured within each 150 μm × 150 μm ROI, using NIS-Elements analysis (Nikon). The mPFC and striatum were identified according to the mouse brain atlas (Franklin and Paxinos, 1997). For quantitative analysis of immunohistochemistry, one ROI per slice × 3 slices in each brain region was used in each mouse.

4. Golgi staining

Golgi staining was carried out with the FD Rapid Golgi Stain Kit (FD NeuroTechnologies, Ellicott City, MD, USA) using the methods described in a previous study[4] . The cryosections were sliced at 80 μm using a cryostat. We obtained the images of layer 2/3 pyramidal neurons in the mPFC by BZ9000 bright-field microscopy (KEYENCE, Osaka, Japan) using a 100×/1.40 NA oil-immersion objective lens. Only fully impregnated neurons isolated from neighboring impregnated neurons were retained for analyses. We quantified the spine density of dendrites 50–200 μm from the soma of pyramidal neurons (six dendrites per mouse). Spine density was expressed as the number of spines per 10 μm of dendrite length. All dendrites and spines were traced in images using Neurolucida software (MicroBrightField Bioscience, Williston, VT, USA) and analyzed using NeuroExplorer (MicroBrightField Bioscience).

5. Behavioral experiments

Touchscreen-based visual discrimination task

The protocol was described in previous reports[4]. Briefly, in order to create enough motivation to perform the task, 8-week-old mice were restricted in their access to food and water to 1 h per day at least 1 week before the pre-training, with the goal of achieving approximately 85–95% of the original bodyweight they attained on an ad libitum diet. The food and water restrictions were continued until the end of the task. The task started with 5 pre-training steps (habituation, initial touch, must touch, must initiate, and punish incorrect) to shape screen-touching behavior. After mice completed this pre-training (≥ 75% correct responses for two consecutive days), they subsequently performed the VD task, in which trial initiation was triggered by their touching the nozzle, and two stimuli (marble and fan) were then presented simultaneously in the two response windows. Touching the correct window (correct response) led to presentation of a liquid reward (20 μL). When the incorrect window was touched (incorrect response), the stimuli offset immediately and a 5-s time-out period was started. After an inter-trial interval (20 s), a correction trial was given instead of a new trial. In the correction trial, the same stimulus set was repeatedly presented in the same location until the mouse made a correct response. Stimulus contingencies were counterbalanced. The session finished after 1 h or 30 trials were completed, whichever came first. When mice could achieve ≥ 80% correct responses for two consecutive days, the final stage, namely the VD task with drug treatment, was begun. In this stage, the initial acquisition and the contingency of the stimulus pair were similar to those of the VD task and METH, MK-801 or KD025 was administered in a crossover design considering the counterbalance. The percentage of correct responses and latency from the correct response to reward retrieval (reward latency) were analyzed.

NORT

The NORT was performed as described previously[3] Mice were individually habituated to an experimental apparatus (30 × 30 × 35 cm) for 10 min every day for three consecutive days (days 1–3). During the training session on day 4, two different objects, different in shape and color but similar in size, were placed symmetrically in the experimental apparatus. Mice were placed individually in the experimental apparatus after KD025 and MK-801 treatment and allowed to explore two objects freely for 10 min, during which the time spent exploring each object was recorded. Mice were readily returned to their home cages after training. During the test session on day 5, one of the familiar objects used during the training was replaced by a novel object. Individual mice were replaced in the same apparatus 24 h after the training session, and the time spent exploring each object was recorded for a 5 min observation period. The preference index in the retention session, the ratio of the amount of time spent exploring the novel object to the total time spent exploring both objects, was used to measure cognitive function.

PPI test

The PPI test was conducted as described previously[3]. After mice were placed individually in the test chamber under 180 lx condition, they were habituated by exposure for 10 min to 65 dB of background white noise. The test consisted of three types of trials: only pulse stimulus (10 startle trials), pulse stimulus with prepulse stimulus (40 prepulse inhibition trials), and only white noise (10 no stimulus trials). The inter-trial interval was 10–20 s, and the entire session lasted 17 min. In the startle trial, a single 40 ms burst of white noise of 120 dB intensity was used. Prepulse inhibition trials consisted of a prepulse (20 ms burst of white noise at an intensity of 69, 73, 77, or 81 dB) that was followed, 100 ms later, by the startle stimulus (120 dB, 40 ms of white noise). Mice were subjected to 60 different trials pseudo-randomly following four different prepulse trials (69, 73, 77, or 81 dB). We verified that each trial was performed 10 times and that no two consecutive trials were identical. After the beginning of the startle stimulus (1 kHz sampling frequency), movement of mice in the startle chamber was measured for 100 ms, and the rectified and amplified data were sent to a computer to calculate the maximum response for 100 ms. The average amplitude of the 10 startle trials was used as the basal startle amplitude. Prepulse inhibition was calculated using the following formula: $100 \times [1-(Prepulse_x/Pulse_{120})]$ %, in which $Prepulse_x$ was the mean of the 10 prepulse inhibition trials ($Prepulse_{69}$, $Prepulse_{73}$, $Prepulse_{75}$, or $Prepulse_{80}$), and $Pulse_{120}$ was the basal startle amplitude.

Spontaneous locomotor test and hyperlocomotion test

The locomotor test was conducted as described in a previous report[3]. Mice were placed individually in a test cage (25 × 30 × 18 cm) and locomotor activity was measured under 15 lx condition using an infrared sensor (NS-DAS-8; Neuroscience, Tokyo, Japan). Spontaneous locomotor activity was measured every 5 min for a total of 180 min after drug treatment. In hyperlocomotion test, mice were habituated in the test cage for 120 min (pre-administration period). Locomotor activity was measured every 5 min for a total of 120 min after METH or MK-801 administration.

Open field test

Open field test was conducted as described in a previous report[9]. Mice were placed at the center of an open field (diameter, 60 cm; height, 35 cm) under a light condition (60 lx) and allowed to explore for 10 minutes while their activity was recorded by using the EthoVision automated tracking program (Noldus, Wageningen, Netherlands). The open field consists of two areas, an inner area (diameter, 40 cm) and an outer area surrounding the inner area. The movement of mice was recorded via a camera mounted above the open field. Measurements of activity included distance traveled and duration in each area. In addition, we manually counted the number of crossing between inner area and outer area, rearing, jumping, grooming, defecation, and urination.

Pole test

The pole test was carried out as previously described, with minor modifications[10, 11]. Briefly, mice were placed head upward at the top of a metal pole (12 mm in diameter and 85 cm in height) wrapped with PROWIPE (Daio Paper Corporation, Tokyo, Japan), and then the times required for the animal to completely rotate downward ($T_{turn}$) and descend to the floor ($T_{total}$) were measured, with a maximum limit of 300 s. The test was repeated two times before and 2, 4, 6, and 24 h after drug treatment, and the average of the trials is reported as the duration. Bradykinesia was defined as the prolongation of $T_{turn}$ and $T_{total}$ values in drug-treated mice compared with vehicle-treated mice. One mouse with a $T_{total}$ over 300 s before treatment and one that preferred to climb the bar were excluded.

Bar test

The bar test was carried out as previously described, with minor modifications[12, 13]. Mice were positioned with their forepaws on a metal bar (9 mm in diameter) 5 cm above the floor, and the time that the mice remained in this position, which was defined as the cataleptic posture, was observed for a maximum of 300 s. The test ended when mice removed themselves from this position by touching the floor with their forepaws or by climbing on the bar. The test was repeated three times before and 2, 4, 6, and 24 h after drug treatment, and the average of the trials is reported as the time on the bar.

6. Measurement of systolic blood pressure and heart rate

Systolic blood pressure and heart rate were measured using a tailcuff system (MK 2000; Muromachi Kikai, Tokyo, Japan), as described in a previous report[14]. Mice were acclimatized to the apparatus by performing daily measurement sessions for 3 days before initiating each experiment. Six blood pressure and heart rate readings were recorded each time, and the average value was used for the analysis. Because of difference in $T_{max}$ of test compounds, we measured blood pressure 20 and 120 min after treatment of fasudil and KD025, respectively. Considering the difference in timing of measurement, we confirmed that blood pressure did not change at all between 20 and 120 min after vehicle treatment (data not shown).

7. Measurement of serum prolactin concentrations

Blood was collected via cardiac puncture and allowed to clot for 30 min at room temperature. Samples were centrifuged at 2000 × *g* for 5 min and serum was collected[15, 16]. Prolactin concentrations were assessed with the Mouse Prolactin ELISA Kit (ab100736; Abcam) using an overnight incubation protocol.

8. Measurement of blood glucose concentrations

Mice were fasted for about 18 h before the experiment[17]. Blood was collected from the tail, and glucose concentrations were measured in duplicate using the FreeStyle Freedom Lite Blood Glucose Monitoring System (11-779; NIPRO CORPORATION, Osaka, Japan).

9. Statistical analysis

All data are expressed as mean + SEM. Statistical analyses were performed with GraphPad Prism 9 (RRID:SCR_002798; GraphPad Software, San Diego, CA, USA). Statistical significance ($P < 0.05$) was determined using One-way analysis of variance (ANOVA), Two-way ANOVA or the Kruskal-Wallis test for multigroup comparisons. Tukey's multiple comparison test, Dunnett's multiple comparisons test, Šídák's multiple comparisons test, or Dunn's multiple comparisons test was used for post hoc comparison. The sample size in each experiment was determined with reference to our previous studies according to experiment[3-5, 10-17]. Detailed information concerning statistical analysis is shown in Supplemental Table S3.
